# Fast qualit**Y** con**T**rol me**T**hod fo**R** der**I**ved diff**U**sion **M**etrics (**YTTRIUM**) in big data analysis: UK Biobank 18608 example

**DOI:** 10.1101/2020.02.17.952697

**Authors:** Ivan I. Maximov, Dennis van der Meer, Ann-Marie de Lange, Tobias Kaufmann, Alexey Shadrin, Oleksandr Frei, Thomas Wolfers, Lars T. Westlye

**Affiliations:** Department of Psychology, University of Oslo, Oslo, Norway; Norwegian Centre for Mental Disorders Research (NORMENT), Department of Mental Health and Addiction, Oslo University Hospital, Oslo, Norway and Institute of Clinical Medicine, University of Oslo, Oslo, Norway; School of Mental Health and Neuroscience, Faculty of Health, Medicine and Life Sciences, Maastricht University, The Netherlands; Department of Psychiatry, University of Oxford, Warneford Hospital, Oxford, UK

**Keywords:** diffusion QC, UK Biobank, DTI, DKI, WMTI, Brain maturation, YTTRIUM

## Abstract

Deriving reliable information about the structural and functional architecture of the brain *in vivo* is critical for the clinical and basic neurosciences. In the new era of large population-based datasets, when multiple brain imaging modalities and contrasts are combined in order to reveal latent brain structural patterns and associations with genetic, demographic and clinical information, automated and stringent quality control (QC) procedures are important. Diffusion magnetic resonance imaging (dMRI) is a fertile imaging technique for probing and visualising brain tissue microstructure *in vivo,* and has been included in most standard imaging protocols in large-scale studies. Due to its sensitivity to subject motion and technical artefacts, automated QC procedures prior to statistical analyses of dMRI data are required to minimise the influence of noise and artefacts. Here, we introduce Fast qualitY conTrol meThod foR derIved diffUsion Metrics (YTTRIUM), a computationally efficient QC method utilising structural similarity to evaluate image quality and mean diffusion metrics. As an example, we applied YTTRIUM in the context of tract-based spatial statistics to assess associations between age and kurtosis imaging and white matter tract integrity in UK Biobank data (n = 18,608). In order to assess the influence of outliers on results obtained using machine learning approaches, we tested the effects of applying YTTRIUM on brain age prediction. We demonstrated that the proposed QC pipeline represents an efficient approach for identifying poor quality datasets and artifacts and increase the accuracy of machine learning based brain age prediction.

## Introduction

Diffusion magnetic resonance imaging (dMRI) provides a range of structural brain features based on routine clinical measurements, which has contributed to its popularity across fields and applications (Kochunov et al. 2015), (de Lange et al. 2019), (Westlye et al. 2010). Advanced dMRI is technically challenging and often involves time-consuming acquisitions placing high demands on the performance and stability of the scanner hardware. Therefore, dMRI data are vulnerable to experimental setup perturbations including post-processing approaches, which might bias the results. In turn, optimised post-processing pipelines (Ades-Aron et al. 2018), (Maximov, Alnæs, and Westlye 2019), (Tournier et al. 2019) and stringent procedures for quality control (QC) (Graham, Drobnjak, and Zhang 2018), (Alfaro-Almagro et al. 2018), (Bastiani et al. 2019) are important to increase reliability and sensitivity. Various approaches have been developed to detect and correct artefacts in raw diffusion data originating, e.g. from eddy currents, bulk head motions, susceptibility distortions (Andersson and Sotiropoulos 2016), noise (Kochunov et al. 2018), presence of outliers (Koch et al. 2019), and diffusion metric variability (Maximov et al. 2015), (David et al. 2019).

However, QC and data harmonisation procedures applied on raw diffusion data (Mirzaalian et al. 2018), (Fortin et al. 2017) do not guarantee accurate numerical computation of scalar diffusion metrics. Derived diffusion metrics from diffusion or kurtosis tensors are sensitive to a range of subject-specific factors such as age or various brain disorders, but also to applied numerical algorithm or its programming implementation (Lebel et al. 2012), (Grinberg et al. 2017), (Maximov et al. 2015), (David et al. 2019). The effects of noisy observations on subsequent between-subjects analysis involving the derived diffusion metrics can be mitigated using simple outlier detection procedures (see, for example, (Richard et al. 2018), (Tønnesen et al. 2018), (de Lange et al. 2019)). However, few publications have directly assessed the effects of QC filtration of final data and performing a sanity check of the derived scalar maps before the statistical analysis. As an example, one can use a visual inspection (see, for example, *slicedir* utility from FSL (Smith et al. 2007)) or truncation based on variability of the data and their standard deviation. We know that outliers might affect the results of analysis, in particular, a machine learning algorithms and related prediction or classification output. An excellent example is an estimation of brain age gap (Smith et al. 2019), (Kaufmann et al. 2019), when a consistency of the used big data sample is vital for an accurate prediction.

Here, we introduce a QC method based on twofold parameterisation: first, diffusion data reduction based on the scalar diffusion values averaged across skeleton voxels using tract-based spatial statistics (TBSS) (Smith et al. 2007), and, second, structural similarity (SSIM) (Wang et al. 2004) of individual diffusion maps relative to the mean diffusion image derived from all subjects. We demonstrate feasibility of this approach for UK Biobank (UKB) data (Miller et al. 2016) using three commonly applied diffusion models: diffusion tensor imaging (DTI) (Basser, Mattiello, and Lebihan 1994), diffusion kurtosis imaging (DKI) (Jensen et al. 2005), and white matter tract integrity (WMTI) (Fieremans, Jensen, and Helpern 2011). We evaluated an effect of the developed QC approach by an analysis of age-diffusion associations along UKB data and variability of the brain age gap estimations using machine learning technique.

## Methods and Materials

### Participants and MRI data

We used dMRI data obtained from 18,608 subjects (see Figure 1 for age and sex distribution). An accurate overview of the UKB imaging acquisition parameters and initial QC pipeline can be found in (Miller et al. 2016), (Alfaro-Almagro et al. 2018). Briefly, a conventional Stejskal-Tanner monopolar spin-echo echo-planar imaging (EPI) sequence was used with multiband factor 3, diffusion weightings (*b*-values) were 1 and 2 ms/µm^2^ and 50 non-coplanar diffusion directions per shell. All subjects were scanned at 3T Siemens Skyra scanners with a standard Siemens 32-channel head coil, in Cheadle and Newcastle, UK. The spatial resolution was 2 mm^3^ isotropic, and 5 AP vs 3 PA images with *b* = 0 ms/µm^2^ were acquired. All diffusion data were post-processed using an optimised diffusion pipeline (Maximov, Alnæs, and Westlye 2019) consisting of 6 steps: noise correction (Veraart et al. 2016), Gibbs-ringing correction (Kellner et al. 2016), estimation of echo-planar imaging distortions, head motions, eddy-current and susceptibility distortions (Andersson and Sotiropoulos 2016), spatial smoothing using *fslmaths* from FSL (Jenkinson et al. 2012) with a 1 mm^3^ Gaussian kernel, and diffusion metrics estimation using Matlab scripts (MathWorks, Natick, MA, USA) (Veraart et al. 2013). UKB data were processed using the high-performance computing facility Colossus at the University of Oslo and large data storage located at Services for Sensitive Data (TSD).

**Figure 1.**
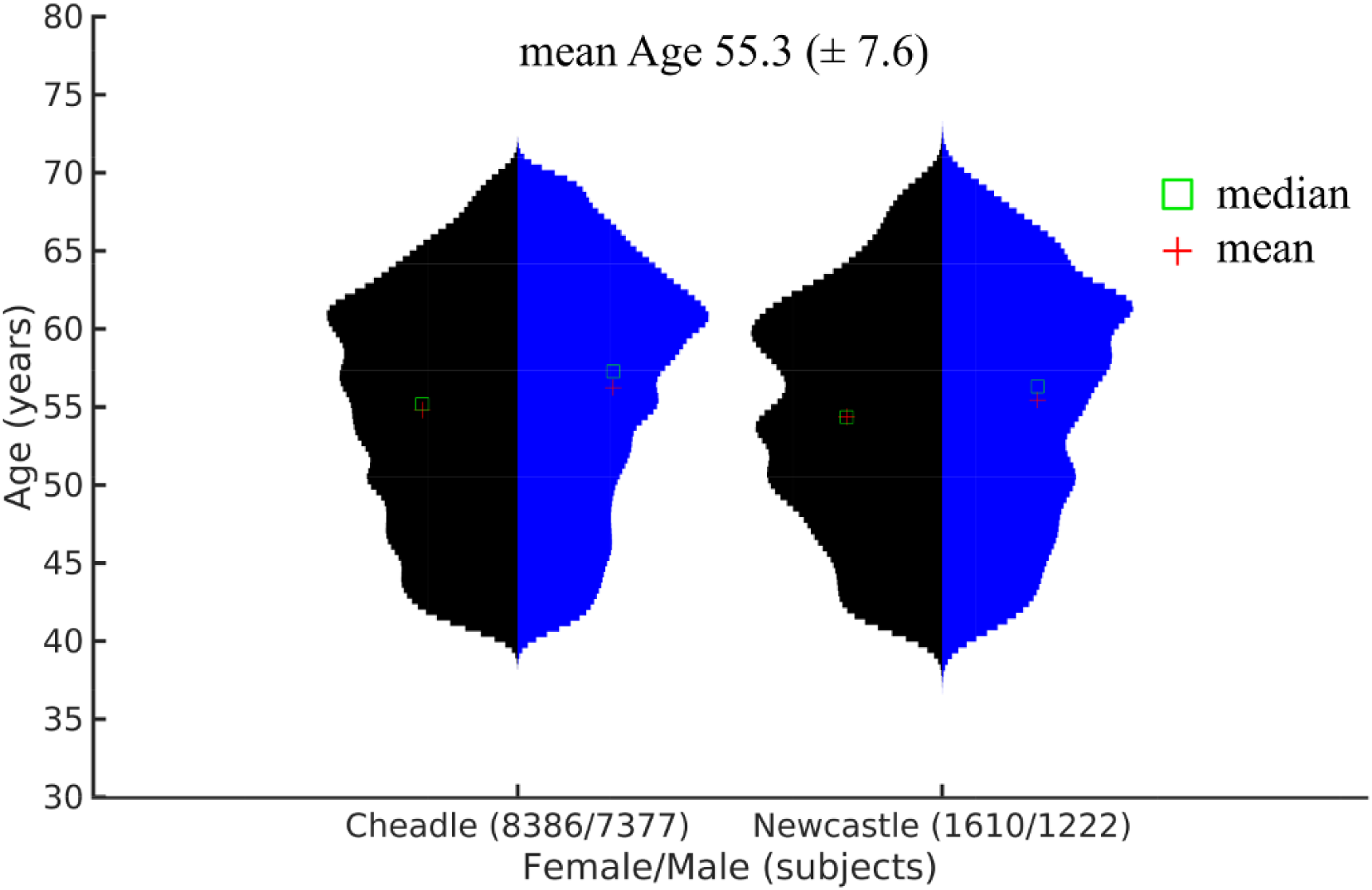
Demographic data depending on the scanner location and gender. The mean age (standard deviation) for all data is on the top of the plot.

### Diffusion metrics

#### DTI and DKI

Diffusion signal decay can be represented as the Taylor expansion along diffusion weightings (Novikov, Kiselev, and Jespersen 2018). This can be approximated by two diffusion tensors of the second (DTI) and fourth (DKI) orders. A set of scalar maps are derived from eigenvalues of the both tensors such as FA, mean, axial and radial diffusivities (MD, AD, RD, respectively), and mean, axial, and radial kurtosis (MK, AK, and RK, respectively). The scalar maps characterise integrative features of brain tissue with potential to represent sensitive biomarkers (Jones 2010).

#### WMTI

In the frame of standard diffusion model (Novikov, Kiselev, and Jespersen 2018), WMTI represents an intra-axonal space as a bundle of cylinders with effective radius equal to zero (Fieremans, Jensen, and Helpern 2011). The cylinders are impermeable, i.e. there is no water exchange between intra- and extra-axonal spaces. The extra-axonal space is described by anisotropic Gaussian diffusion. In order to keep the model simple a few more assumptions have been made: intra-axonal space consists of mostly myelinated axons without any contribution from myelin due to a fast relaxation rate across of typical diffusion times; at the same time in extra-axonal space the glial cells possess fast water exchange with extra-cellular matrix; both intra- and extra-axonal spaces are modelled by Gaussian diffusion tensors. In order to avoid degeneration (Jelescu et al. 2016), intra-axonal diffusion is assumed to be slower than diffusion in extra-axonal matrix. Besides, WMTI parametrisation works in the case of quite coherent axonal bundle with an orientation dispersion below 30° (Fieremans, Jensen, and Helpern 2011). WMTI allows one to derive axonal water fraction (AWF), extra-axonal axial and radial diffusivities (axEAD and radEAD, respectively).

### Tract Based Spatial Statistics

In order to evaluate and compare different QC approaches, we applied TBSS (Smith et al. 2007). Initially, all FA volumes were aligned to the FMRI58_FA template, supplied by FSL, using non-linear transformation implemented by FNIRT (Andersson and Jenkinson 2007). Next, a mean FA image across 18,600 subjects was obtained and thinned in order to create mean FA skeleton. Afterwards, each subject’s FA data are projected onto the mean FA skeleton, by filling the skeleton with FA values from the nearest relevant tract centre. TBSS minimises confounding effects due to partial voluming and residual misalignments originated from non-linear spatial transformations. For each diffusion metric we computed the individual skeleton projecting the non-FA values onto the FA skeleton.

### QC Model description

Our approach of image quality estimation originates from multidimensional experiments in nuclear magnetic resonance spectroscopy (Ernst, Bodenhausen, and Wokaun 1987), when an additional dimension allows one to resolve hidden resonance peaks. Natural parameter in diffusion scalar metrics is an absolute value which either has its physical limitations, for example, as FA lies between 0 and 1, or some region-specific values from other sources, for example, a free water diffusivity in the brain equals to 3 μm^2^/ms. Thus, by applying a reasonable threshold rule one can discard volumes with unfeasible values from further analysis. However, since the values are typically averaged over the volume (region of interest, skeleton etc), we still can expect “hidden” outliers with minimal influence on the averaged metric. We assume that SSIM allows us to spread image parameterisation into the second dimension using three principal features: luminance, contrast, and structure as following (Wang et al. 2004),

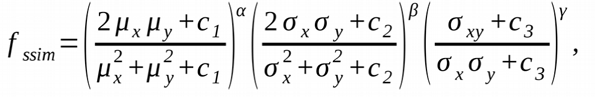

where µ_x,y_ are the mean of *x* and *y*, σ^2^_x,y_ are the variance of *x* and *y*, and σ_xy_ is the covariance of *x* and *y*, constants c_1,2,3_ are the variables stabilising the SSIM estimation, and α, β, and γ are the weights of three SSIM features. In theory, SSIM estimator can be improved for some specific purposes (Charrier et al. 2012) or generalised (Brunet, Vrscay, and Zhou Wang 2012). Nevertheless, original SSIM metric already proved its capability in medical image quality verification (Renieblas et al. 2017), (Chow and Paramesran 2016), (Vinding et al. 2017). The weights α, β, and γ allow one to emphasise the principle SSIM features in order to enhance a contrast between original image and reference. While SSIM weights adjustments is still debated (Li and Bovik 2009), we empirically define the following weights α = 0.1, β = 0.1, and γ = 2, in order to stretch a range of SSIM value components.

In order to verify our approach for detection of possible “badly” estimated scalar metrics and outliers we used a random subset of UKB data consisting of 724 subjects. Next, we manually introduced three types of image distortions. The first type (Type 1) is based on complete loss of *N*_1_ random slices in the image volume. In our case, we set upper bound of *N*_1_ to be equal 5. The second type of artefacts (Type 2) is based on value scaling of up to *N*_2_ = 7 random slices. Scaling of the random slices can be performed in two ways as a division or multiplication of the diffusion values. To dilute scaled values between neighbouring slices we applied 3D Gaussian smoothing with 3mm^3^ kernel. The final type of distortions (Type 3) is based on residual misalignments along an image normalisation process. As a simple implementation of the residual misalignments we used rotation around superior-inferior axis with a random angle up to 5 degrees. An example of original image and three types of artificial distortions is presented in Figure 2. Finally, we added 25 volumes (10.4% of original data) of each type of distortions to four diffusion metrics: FA, MD, MK, and AWF. In order to test influence of artefacts on the derived mean metric maps, we evaluated SSIM parameters of initial datasets using mean maps with (799 volumes) and without (724 volumes) outliers. Resulting SSIM correlations are presented in Fig. 2. High linear correlations (over 0.999) demonstrated that an introduction of outliers into data subsets did not influence on mean reference images and, consequently, on the SSIM evaluations.

**Figure 2.**
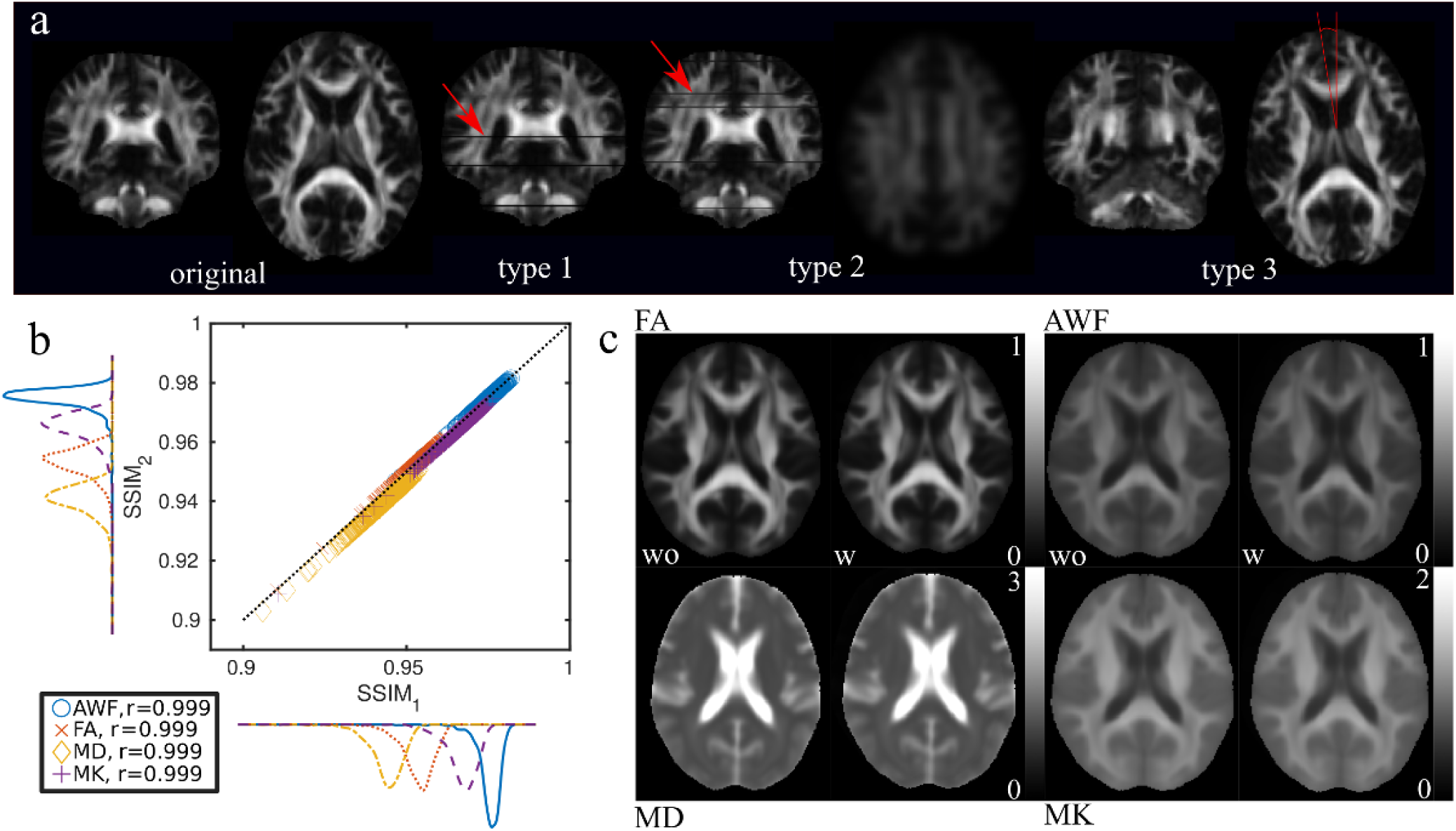
Example of simulated distortions and their influence on the estimated SSIM values. **a)** an original FA image and three types of distortions: type 1 – random zero slices (see the red arrow), type 2 – slices with scaled and smoothed values (see the red arrow), type 3 – rotation of a whole volume around the superior-inferior axis (red lines show the angle); **b)** correlation of the SSIM metrics evaluated for FA, AWF, MD, and MK metrics using data without (SSIM_1_) and with (SSIM_2_) outliers. *r* is the Pearson correlation coefficient; black dotted line is a unity line; **c)** left-side images are mean metrics averaged without (wo) outliers, when right-side images are mean metrics including (w) the outliers into averaging step.

We performed outlier detection using the following two-step approach: first, we used *k*-means clustering (implemented as *kmeans* Matlab function) to define one cluster based on squared Eucledian distances (Arthur and Vassilvitskii 2007). Next, in order to introduce object density parametrisation, we used the median distance of the distance distribution (MDD) around the cluster centroid as a unite for a neighbourhood radius in density-based scan algorithm with noise (*dbscan*, implemented as Matlab function) (Ester et al. 1996), (Daszykowski, Walczak, and Massart 2002). The optional parameters in the *dbscan* algorithm are number of objects in neighbourhood of a central object and number of MDD units. We empirically set it to be equal to 10 and 7, respectively, in the tests. In such a way chosen parameters allow us to apply cluster density estimation independently from the original data distribution (see, for example, Figure 3 and Kendall correlation coefficients between diffusion metrics and SSIM values, estimated using Matlab function *corr*).

**Figure 3.**
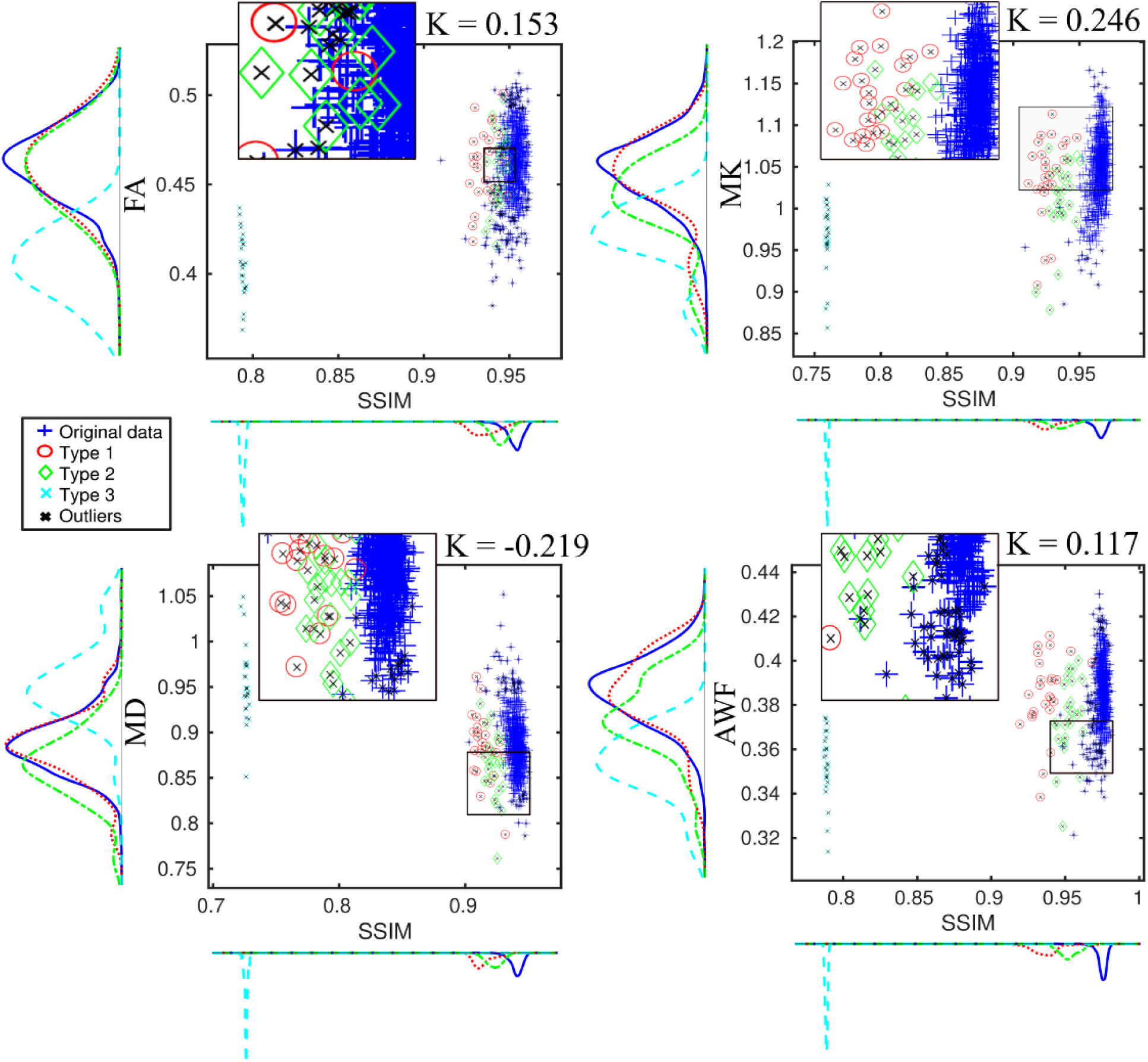
Scatter plots of diffusion metrics and SSIM values for the data with three types of outliers. All arteficial outliers are marked by the different colours. The result of QC method is marked by black crosses. Image inserts demonstrate a zoomed boundary between outlier groups and original data. Estimated Kendall correlation coefficients (K) for the diffusion metrics and SSIM are presented as well. SSIM, FA, MK, and AWF are unit-less values, MD is in µm^2^/ms.

As a frequently applied QC approach for a comparison purpose, we applied a simple threshold approach of |3| standard deviations from the mean diffusion value after regressing out main effects of age, sex and site.

### Statistical analysis

In order to assess the effects of our proposed QC pipeline on the sensitivity of the diffusion metrics we tested for associations with age and sex using linear models as implemented in the Matlab function *lmfit*. In the subsample of 799 subjects (724 UKB subjects + 75 simulated images) we employed the following general linear model (GLM): *y* = *b*_0_ + *b*_1_ Age + *b*_2_ Sex + *b*_3_ Site, where Age is given in years, Sex and Site as a dichotomous variable. We computed specificity and sensitivity of the automated artefact correction. Sensitivity is defined as a ratio of True Positive/(True Positive + False Negative)) and specificity as a ratio of True Negative/(True Negative + False Positive)).

In the full sample we employed the following models: *y* = *b*_0_ + *b*_1_ Age + *b*_2_ Sex + *b*_3_ Site + (*b*_4_ Age^2^). We computed root mean squared error and R-squared as proxies for goodness-of-fit. We compared coefficients between models (before and after discarding datasets flagged by our QC pipeline) using the R package *cocor* (Diedenhofen and Musch 2015). In order to assess normality of the residuals from the linear models we used QQ-plots (Aldor-Noiman et al. 2013) (implemented as *qqplot* Matlab function) and Kolmogorov-Smirnov (KS) test with W critical value (Kolmogoroff 1933), (Smirnov 1948). The W critical values have been used as indirect measures of normality of the residuals. The KS tests were implemented as Matlab function *kstest*.

### Machine learning for brain age gap estimation

We estimated the influence of outliers on machine learning (ML) based brain age prediction. The brain age gap (BAG) is defined as the difference between chronological and predicted age and has been proposed to reflect a sensitive imaging-derived phenotypes (Kaufmann et al. 2019). For age prediction we applied two frequently used approaches. First, we employed linear model and multiple regressors (LMMR) defined as Y = Xβ – δ, where Y is the chronological age, β is the regressor vector, X is the matrix of brain features used for prediction, and δ is the BAG. The solution can be obtained by pseudo-inversion X^+^ matrix (Smith et al. 2019). In order to improve the ML-training, we used 25% of eigenstates produced by the singular value decomposition replacing the X matrix as recommended by Smith and colleagues, 2019. The estimations were performed using original Matlab script from Smith et al., 2019 (http://www.fmrib.ox.ac.uk/BrainAgeDelta) with removed cross-validation code. The second algorithm is XGBoost employing a gradient boosting approach (Chen and Guestrin 2016). The parameters of XGBoost were chosen as follows: eta = 1; number of rounds = 250; max depth = 4; lambda = 10^-7^. Estimations were performed using Julia (https://julialang.org) implementation of XGBoost algorithm (https://github.com/dmlc/XGBoost.jl).

For simplicity, we used the following linear model using only 4 diffusion metrics averaged over the skeleton: Y ∼ *b*_1_ FA + *b*_2_ MD + *b*_3_ MK + *b*_4_ AWF + *b*_5_ Sex + *b*_6_ Site. In order to assess the influence of outliers on age prediction, a fixed number of 476 outliers, identified by the proposed QC approach over all diffusion metrics, was combined with varying samples of good-quality data, creating total training-sets of 1000, 2000, 3000, 4000, 5000, 7500, 10000, 12500, and 15000 subjects. The 476 outliers were manually added to each sample, leading to outlier percentages of 47.6%, 23.8%, 15.9%, 11.9%, 9.52%, 6.35%, 4.76%, 3.81%, and 3.17%, respectively. In all training sets we kept the sex and site distribution identical. All training sets were selected from the whole UKB dataset. 1000 subjects not included in the training sets were selected as a test sample that was used in all runs. We performed the BAG estimations separately for the training sets with and without outliers, respectively. As criteria we used the Pearson correlations between chronological and predicted ages, and root mean squared errors estimated for the test sample.

## Results

### Sensitivity to simulated artefacts

Figure 3 shows an application of the developed QC method to the subsample consisting of 799 subjects with artificially introduced outliers of the three types. Briefly, based on AWF we detected all three types of introduced outliers and discarded 112 volumes of original data. The sensitivity of the QC method to all three types of outliers was 1, specificity was 0.85. In the case of FA we missed 10 (13%) outliers (2 of type 1 and 8 of type 2) and discarded 142 volumes of original data (sensitivity = 0.92, 0.68, 1 for Types 1,2,3, respectively; specificity = 0.80); in the case of MD we missed 7 (9%) outliers (1 of type 1 and 6 of type 2) and discarded 69 volumes of original data (sensitivity = 0.96, 0.76, 1; specificity = 0.91); in the case of MK we missed 2 (3%) outliers (type 2) and discarded 21 volumes of original data (sensitivity = 1, 0.92, 1; specificity = 0.97).

Figure 4 shows the various diffusion metrics plotted as a function of age and the corresponding linear fits based on the GLMs. Briefly, the raw data and thresholding method yielded similar GLM parameters (see Table 1 for the intercept and slope values). Cocor revealed no significant slope differences for any of the diffusion metrics between the raw and QC’ed data. Table 1 summarises the goodness-of-fit measures for the selected diffusion metrics for three datasets (raw data, thresholding QC and the developed QC method) and GLM parameters (*b*_0_, intercept, and *b*_1_, age slope). QQ plots and the W parameters from KS tests based on FA, MD, MK, and AWF for three datasets (raw data, thresholding QC, and our QC method) suggested that our proposed QC method yields the most “normal” residuals (Figure 5).

**Figure 4.**
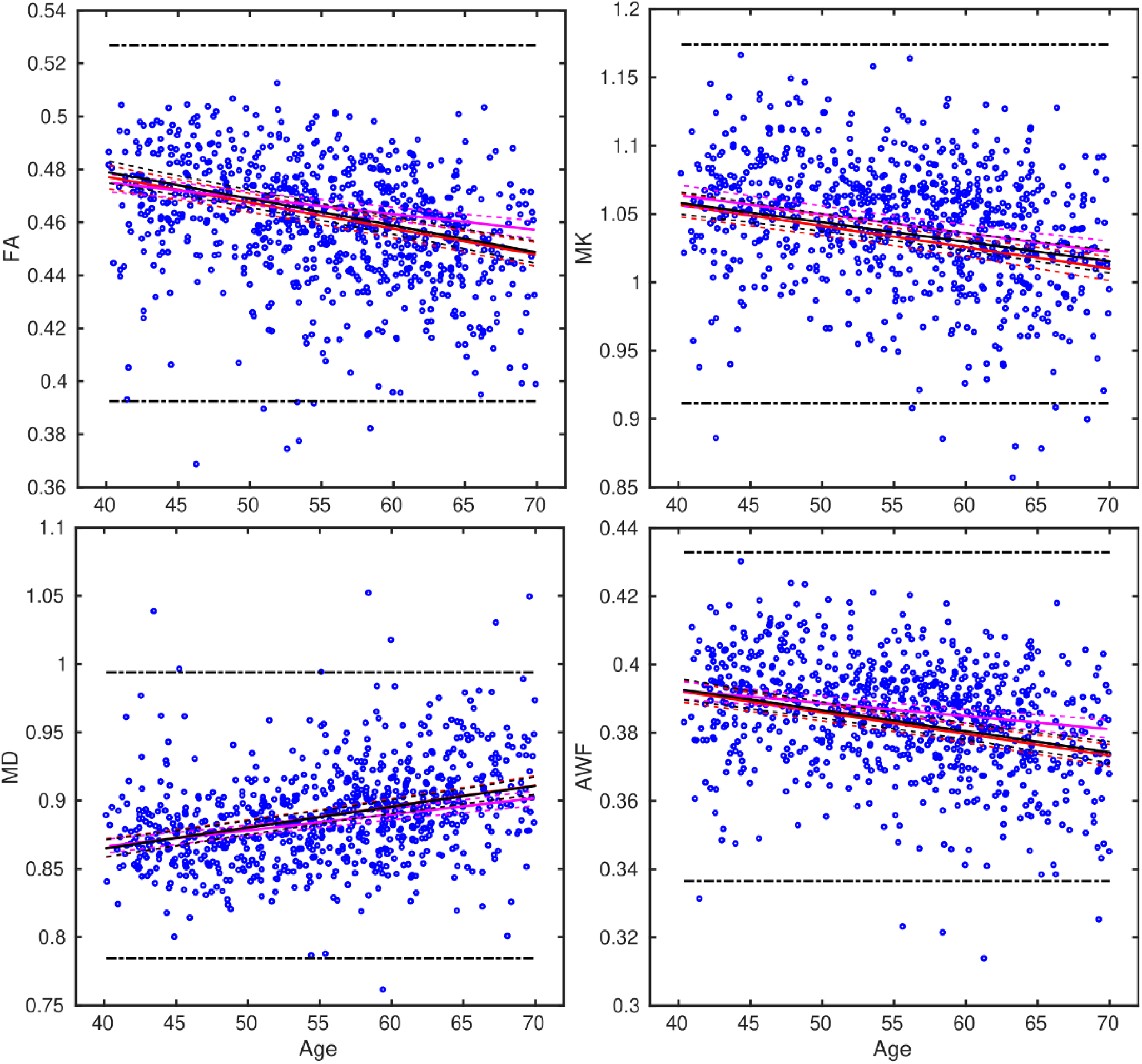
Results of general linear model *y* = *b*_0_ + *b*_1_ Age + *b*_2_ Sex + *b*_3_ Site for 4 diffusion metrics over the test data sample of 724 subjects. The solid and dashed red lines are linear fit (LF) and the interval of confidence (CI 95%); the black solid and dashed lines are LF and CI for data thresholded by 3 standard deviations from the mean diffusion metrics (marked as dot-dash lines); the margenta solid and dashed lines are LF and CI for the proposed QC method. SSIM, FA, MK, and AWF are unit-less values, MD is in µm^2^/ms.

**Figure 5.**
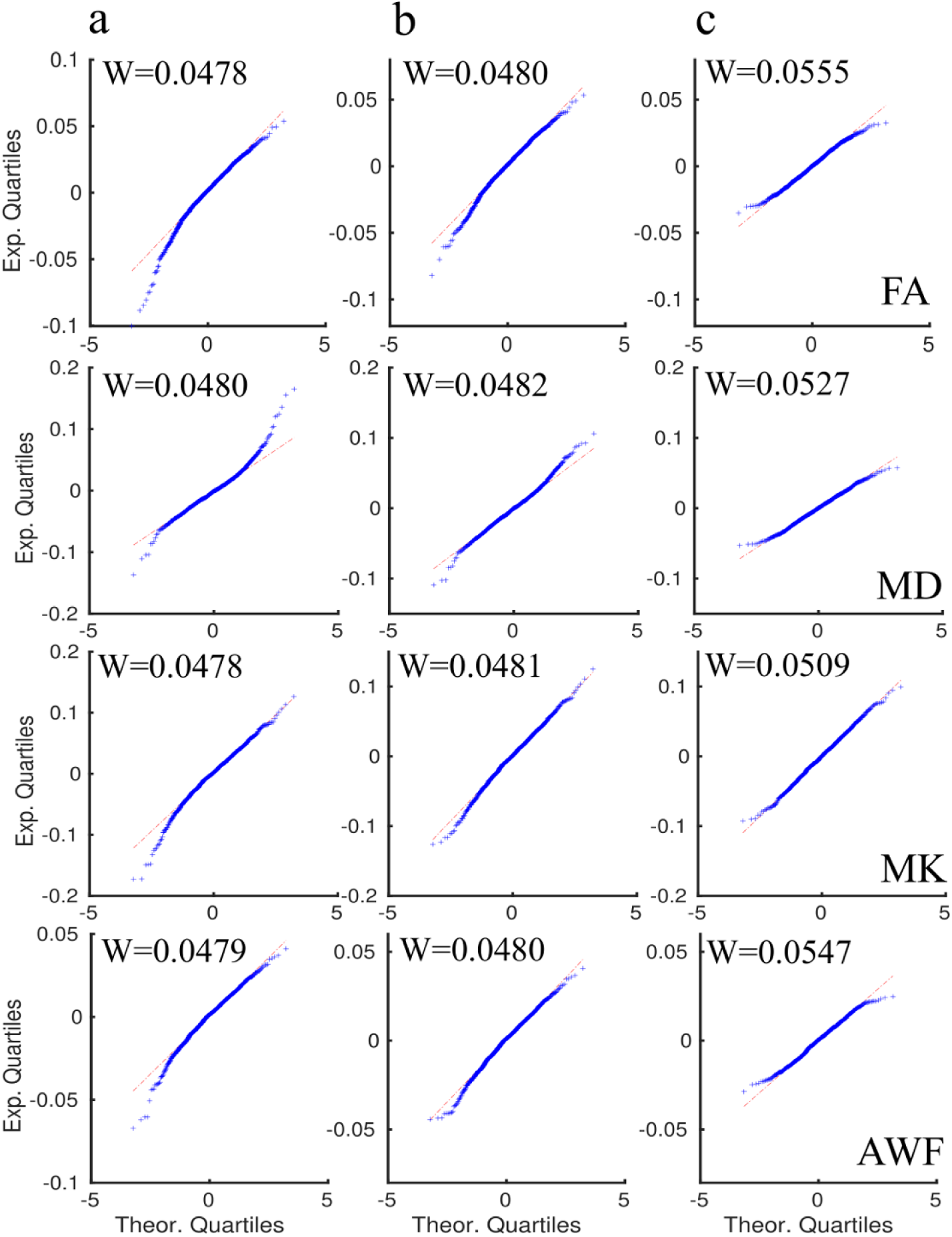
QQ plots of the GLM residuals for three cases (see Fig.4): a) raw data; b) after thresholding by 3 standard deviations; c) the proposed QC method. Values W are the critical numbers of Kolmogorov-Smirnov (KS) test for a normality. In all cases KS test did not reveal that the residuals are normally distributed.

**Table 1.** Results of GLM *y* = *b*_0_ + *b*_1_ Age + *b*_2_ Sex + *b*_3_ Site for four diffusion metrics using test sample of 724 subjects. RMSE is the root mean squared error; R^2^ is the R-squared parameter; NO is the number of observations; STD is the standard deviations; Intercept is *b*_0_; Slope is *b*_1_;

| Metrics/<br>statistics | Raw data<br>NO 799 |  | Threshold with 3 std |  |  | Our QC method |  |  |
| --- | --- | --- | --- | --- | --- | --- | --- | --- |
| | RMSE | $R^2$ | NO | RMSE | $R^2$ | NO | RMSE | $R^2$ |
| **FA** | 0.0211 | 0.112 | 792 | 0.0198 | 0.137 | 592 | 0.0135 | 0.1 |
| **MD** | 0.0327 | 0.125 | 786 | 0.03 | 0.141 | 657 | 0.0223 | 0.144 |
| **MK** | 0.0416 | 0.0999 | 791 | 0.0391 | 0.0985 | 705 | 0.0345 | 0.105 |
| **AWF** | 0.0152 | 0.103 | 793 | 0.0145 | 0.108 | 611 | 0.0109 | 0.0629 |
|  | **Intercept** | **Slope** |  | **Intercept** | **Slope** |  | **Intercept** | **Slope** |
| **FA** | 0.5164 | $-9.80 \cdot 10^{-4}$ | | 0.5197 | $-1.01 \cdot 10^{-3}$ | | 0.4990 | $-5.98 \cdot 10^{-4}$ |
| **MD** | 0.8037 | $1.54 \cdot 10^{-3}$ | | 0.8037 | $1.53 \cdot 10^{-3}$ | | 0.8194 | $1.18 \cdot 10^{-3}$ |
| **MK** | 1.1188 | $-1.55 \cdot 10^{-3}$ | | 1.1154 | $-1.43 \cdot 10^{-3}$ | | 1.1189 | $-1.38 \cdot 10^{-3}$ |
| **AWF** | 0.4174 | $-6.29 \cdot 10^{-4}$ | | 0.4176 | $-6.19 \cdot 10^{-4}$ | | 0.4077 | $-3.80 \cdot 10^{-4}$ |

### Effects of QC pipeline on the sensitivity to age, sex and scanner site

Figure 6 shows an application of the developed QC method to the UKB data with 18,608 subjects. As detailed above, we discarded datasets defined as outliers based on mean skeleton diffusion metrics and SSIM. The mean diffusion maps used as a reference for SSIM estimations are depicted in Supplementary Materials. Distributions of relevant diffusion metrics and demographics of the data defined as outliers are presented in Supplementary Materials. A higher number of outliers were identified from the Cheadle (n = 396, 3%) site compared to the Newcastle (n = 78, 1%) site, and both sex (39% of women) and age distribution (58.55/7.74 years) of the outlier data did not diverge substantially from the distributions in the total sample.

**Figure 6.**
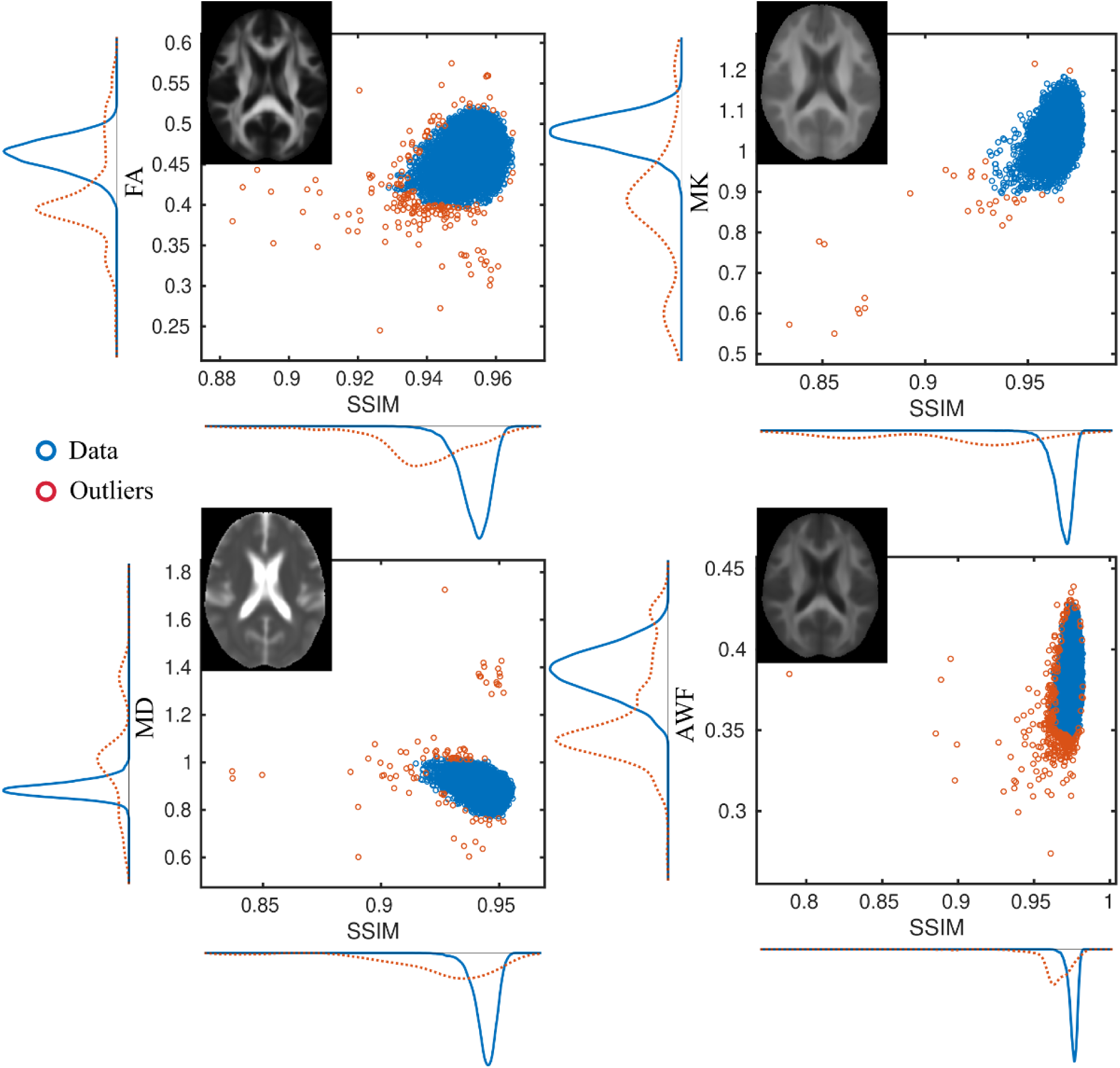
An application of QC method to UKB data. Mean diffusion maps are presented as a reference for the SSIM estimations. The red circles are identified outliers; the blue circles are the filtered data. SSIM, FA, MK, and AWF are unit-less values, MD is in µm^2^/ms.

Figure 7 presents age-related trajectories (linear and quadratic fits) for four diffusion metrics with the detected outliers included or excluded. The summary statistics are summarised in Table 2. All other metrics are shown in the Supplementary Materials. Figure 8 shows corresponding QQ-plots of the GLM residuals in the case of linear and quadratic age terms. QQ-plots of the residuals for all diffusion metrics are shown in Supplementary Materials. Briefly, the residuals from the models using raw data show strong deviations from the diagonal in both linear and quadratic age terms. In contrast, the residuals from QC approved data appear more normal.

**Figure 7.**
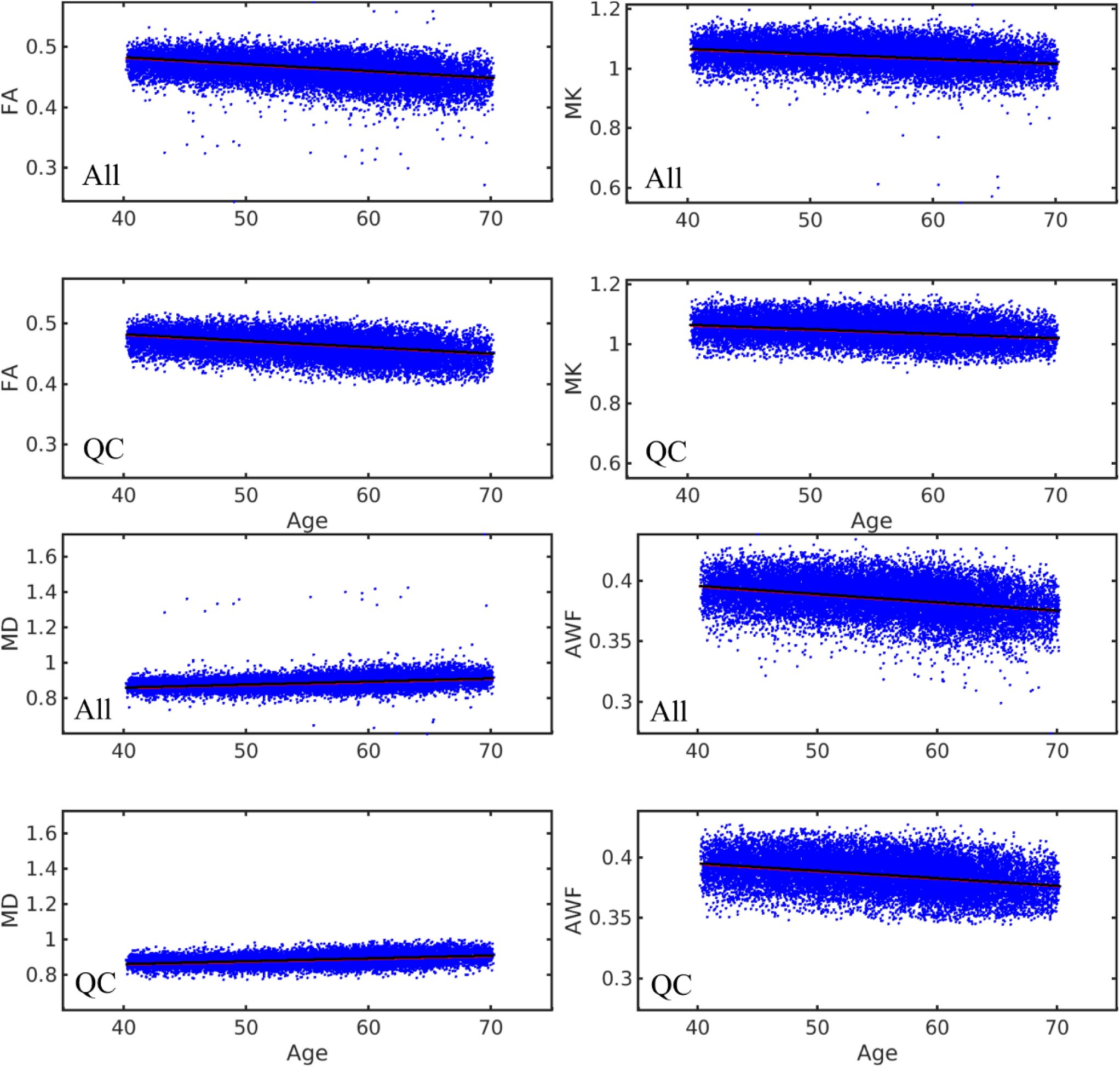
The results of GLM age-diffusion correlations with linear age term (red line) and quadratic age term (black line). The plots marked as “All” consists of all raw data (n = 18608); the plots marked as “QC” consists of data passed through the QC filtration (n = 18132). Intervals of confidence (CI 95%) are presented as dashed line in all cases. FA, MK, and AWF are unit-less values, MD is in µm^2^/ms.

**Figure 8.**
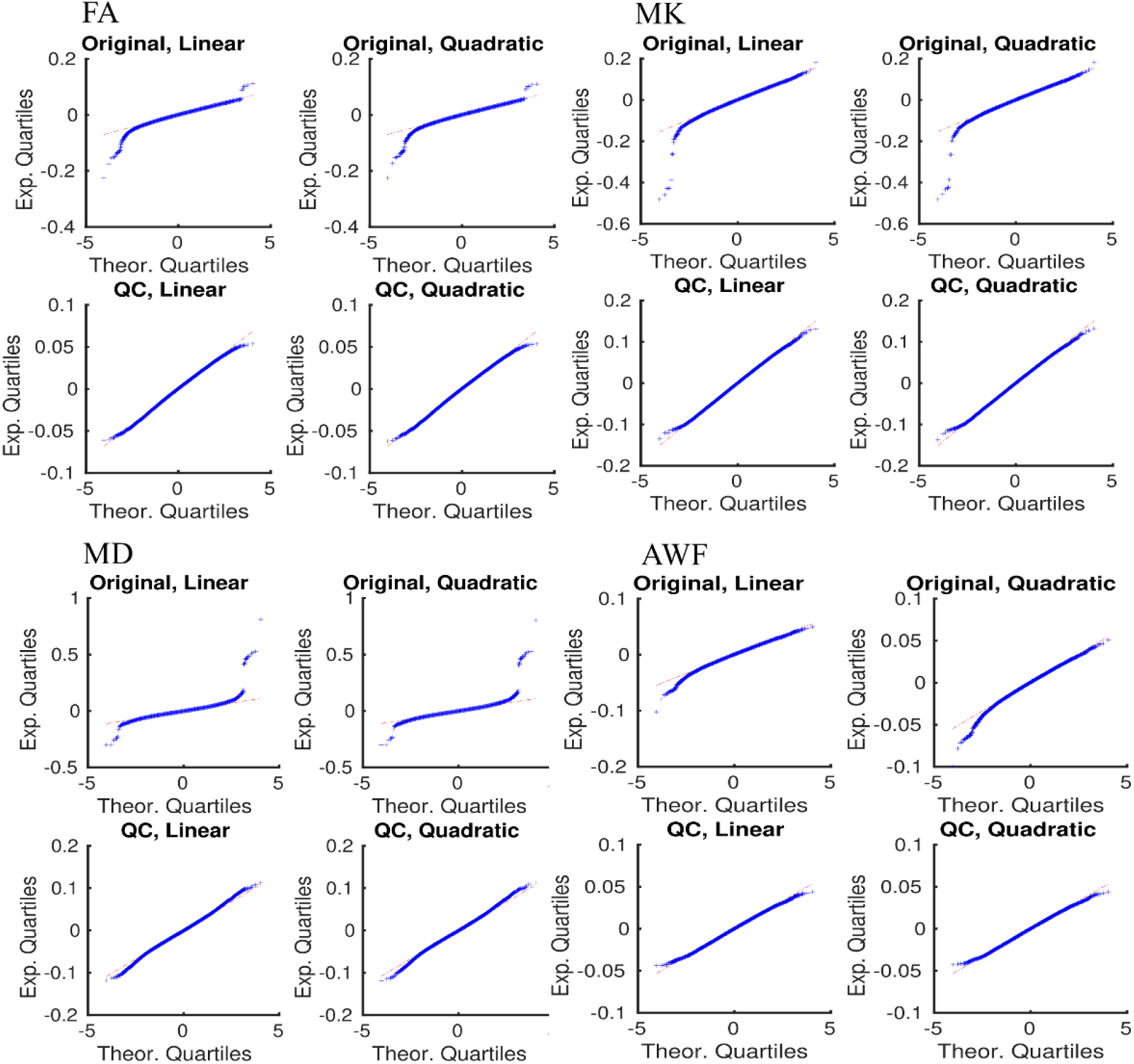
QQ-plots of the GLM residuals for four diffusion metrics. “Original, Linear” means the all data and GLM with linear age term; “Original, Quadratic” means the all data and GLM with quadratic age term; “QC, Linear” means the QC filtered data and GLM with linear age term; “QC, Quadratic” means the QC filtered data and GLM with quadratic age term.

**Table 2.** Results of two GLM *y* = *b*_0_ + *b*_1_ Age + *b*_2_ Sex + *b*_3_ Site + (*b*_4_ Age^2^) for four diffusion metrics using all UKB data with and without QC procedure. RMSE is the root mean squared error; R ^2^ is the R-squared parameter, NO is the number of observations; Intercept is *b*_0_; Slope is *b*_1_.

| Metrics/<br>statistics | All data, linear Age term |  |  | QC passed, linear Age term |  |  |
| --- | --- | --- | --- | --- | --- | --- |
|  | NO | RMSE | R<sup>2</sup> | NO | RMSE | R<sup>2</sup> |
| FA | 18608 | 0.0187 | 0.173 | 18134 | 0.0171 | 0.179 |
| MD | 18603 | 0.0337 | 0.133 | 18129 | 0.0282 | 0.158 |
| MK | 18608 | 0.0398 | 0.1 | 18134 | 0037 | 0.096 |
| AWF | 18607 | 0.014 | 0.119 | 18133 | 0.0131 | 0.113 |
|  |  | Intercept | Slope |  | Intercept | Slope |
| FA |  | 0.5277 | -1.12·10<sup>-3</sup> |  | 0.5242 | -1.05·10<sup>-3</sup> |
| MD |  | 0.7914 | 1.70·10<sup>-3</sup> |  | 0.7976 | 1.58·10<sup>-3</sup> |
| MK |  | 1.1315 | -1.64·10<sup>-3</sup> |  | 1.1231 | -1.47·10<sup>-3</sup> |
| AWF |  | 0.4226 | -6.70·10<sup>-4</sup> |  | 0.4196 | -6.10·10<sup>-4</sup> |
|  | All data, quadratic Age term |  |  | QC passed, quadratic Age term |  |  |
|  | NO | RMSE | R<sup>2</sup> | NO | RMSE | R<sup>2</sup> |
| FA | 18608 | 0.0187 | 0.176 | 18134 | 0.0171 | 0.182 |
| MD | 18603 | 0.0335 | 0.141 | 18129 | 0.028 | 0.165 |
| MK | 18608 | 0.0397 | 0.106 | 18134 | 0.0369 | 0.101 |
| AWF | 18607 | 0.014 | 0.125 | 18133 | 0.0131 | 0.117 |
|  |  | Intercept | Slope |  | Intercept | Slope |
| FA |  | 0.5283 | -1.13·10<sup>-3</sup> |  | 0.5247 | -1.06·10<sup>-3</sup> |
| MD |  | 0.7898 | 1.73·10<sup>-3</sup> |  | 0.7962 | 1.61·10<sup>-3</sup> |
| MK |  | 1.1332 | 1.67·10<sup>-3</sup> |  | 1.1246 | -1.50·10<sup>-3</sup> |
| AWF |  | 0.4232 | -6.81·10<sup>-4</sup> |  | 0.4201 | -6.19·10<sup>-4</sup> |

### Effects of QC pipeline on the ML BAG estimations

Figure 9 shows the trajectories of RMSE and correlations between chronological and predicted ages for the two ML algorithms. For statistics and cross validation of the BAG results, we repeated model training 100 times, randomly choosing the training samples from whole UKB data. Briefly, in the case of QC filtered data model performance increased only moderately with sample size, with RMSE of the XGBoost algorithm suggesting only minor effects. In contrast, the training sets with outliers demonstrated strong dependence of the chronological and predicted age correlations and RMSE on the percentage of outliers in the training sample, in particular, for the LMMR approach. For both ML algorithms, increasing training sample size decreased RMSE in the test set.

**Figure 9.**
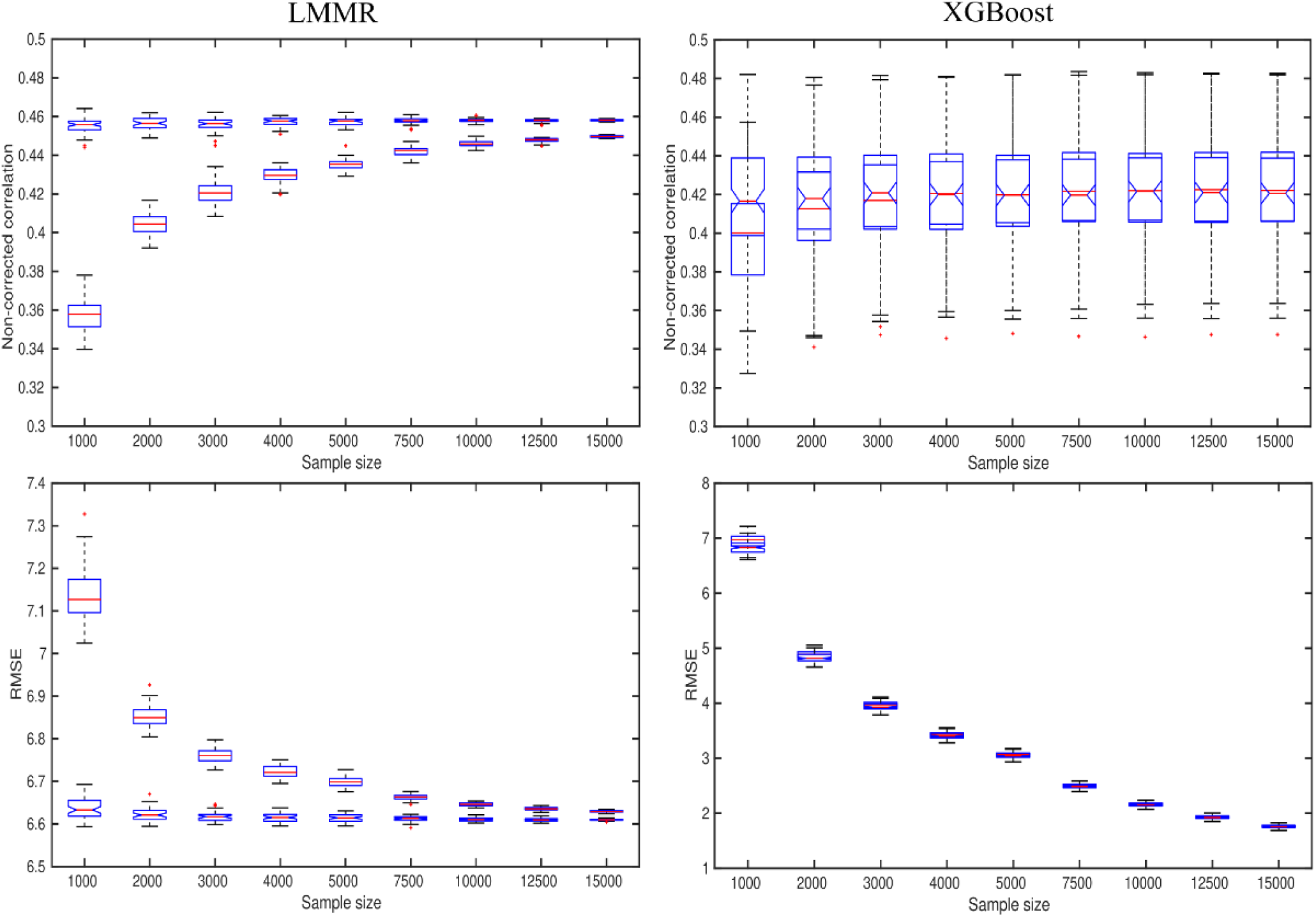
Outlier influence on the brain age predictions for two ML algorithm: linear model with multiple regressors (LMMR) and gradient boosting method (XGBoost). The top row is non-corrected Pearson correlations between chronological and predicted ages as a function of the sample size of training sets. The correlations were estimated for the fixed test sample of 1000 subjects. The bottom row is root mean square error (RMSE) of the predictions as a function of the sample size of training sets. The rectangular boxplots are the datasets with included outliers, the boxplots with “notched” feature are the datasets without outliers (QC filtered).

## Discussion

Advanced dMRI offers sensitive measures of brain tissue micro- and macrostructural architecture and integrity, with large potential for the basic and clinical neurosciences. With the surge of large-scale clinical and population-based efforts acquiring dMRI data from thousands of individuals, there is an increasing need to develop computationally efficient pipelines for quality assessment and identification of poor quality data. The proposed QC method based on 2D data representation exploiting similarity metrics and data density features enables an efficient evaluation of data quality after estimation of diffusion scalar metrics. In a subsample (n = 799), our semi-automated artefact detection based on similarity metrics yielded high sensitivity and specificity and the residuals from linear age fits in the full sample (n = 18,608) resulted in more normal residuals after compared to before discarding flagged datasets. Additionally, the QC pipeline improved brain-age prediction using machine learning by mitigating the influence of outliers in the training set.

By default, the harmonised validated raw diffusion data allow one to derive accurate scalar metrics. However, a quality evaluation of the processed diffusion maps is still an open question in big data analysis. Many efforts have been made to develop accurate QC and harmonisation procedures on raw diffusion weighted data (Mirzaalian et al. 2018), (Fortin et al. 2017). Nevertheless, derived diffusion metrics from DTI or DKI may still deviate from expected range, e.g. due to remaining artefacts and numerical misestimation. The simple considerations of 2D representations of the averaged diffusion metric and SSIM values is an advantage of the developed QC method. This allows us to take into account frequently applied measures in large-scale studies, i.e. diffusion metrics averaged across a region of interest or the entire TBSS skeleton and the structural similarity based on the intensity, contrast, and structure of the scalar map in relation to a reference maps. Our simulations revealed that our method is capable of identifying image artefacts with different origins with high sensitivity and specificity. This is particularly valuable in the context of large-scale studies, where manual QC is not feasible and when a quantitative estimate of structural similarity is needed. Whereas our direct comparison of slopes did not reveal a significant effect of the QC procedure on the estimated age-associations, the linear models based on QC’ed data yielded evidence of improved model fits in terms of the distributions of the model residuals compared to models based on the non-QC’ed data.

We found evidence of improved machine learning based age prediction when limiting the number of noisy datasets in the training set. In general, larger training sets are expected to increase accuracy of brain age prediction (see e.g. (Kaufmann et al. 2019)). However, in practice, the number of accessible data is usually limited. Thus, it is very important to know how different amounts of undetected outliers in the training set could affect the prediction accuracy in an independent test set. Our results demonstrated that a higher portion of outliers in the training set influenced the prediction accuracy in the test set. Surprisingly, for XGBoost, in contrast to the RMSE, the correlation between predicted and chronological age did not increase with increasing training set size. For LMMR the correlation coefficients increased in accordance with increased sample size. In both instances, however, the proportion of bad datasets in the training set influenced the prediction accuracy in the test set, with markedly improved prediction with lower proportion of noisy data.

In summary, although an overall beneficial effect of removing poor quality datasets results is not surprising, our results serve as relevant demonstrations of the importance of QC in the context of large-scale studies. It should also be noted that all datasets included in the current analysis have been checked and approved by the initial UK Biobank QC procedures (Alfaro-Almagro et al. 2018), and the reported effects of noise removal on age-associations are likely to represent lower-bound effects compared to a scenario with no initial QC procedures. In general, whereas minimizing noise is a universal aim, the direct effects and value of QC will vary between studies and applications.

Conclusively, in the case of big data, automated, efficient and reliable approaches for evaluating the scalar diffusion metrics prior to statistical analysis are needed. Our results suggest that our proposed method is suitable as a complementary test of the estimated diffusion data to increase sensitivity of conventional diffusion scalar metrics.

## Acknowledgement

This work was funded by the Research Council of Norway (249795). This research has been conducted using the UK Biobank under Application 27412. IIM thanks Dr. Viljami Sairanen for his help with FSL and *slurm* queue scripting.

## 1. Mean diffusion maps averaged over evaluated subjects (18608) in MNI space

Cross section coordinates for all diffusion metrics: X = 75; Y = 108; Z = 78

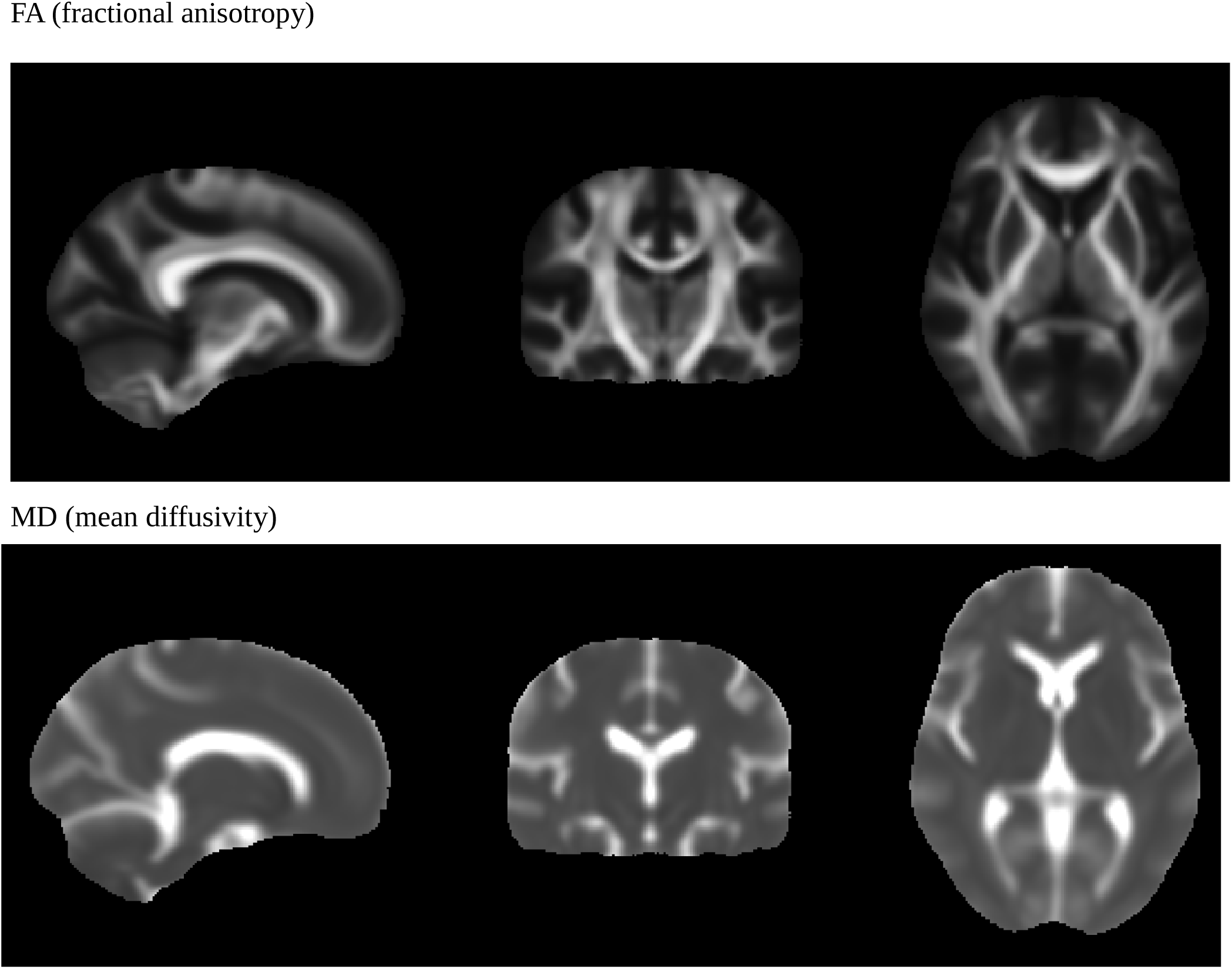

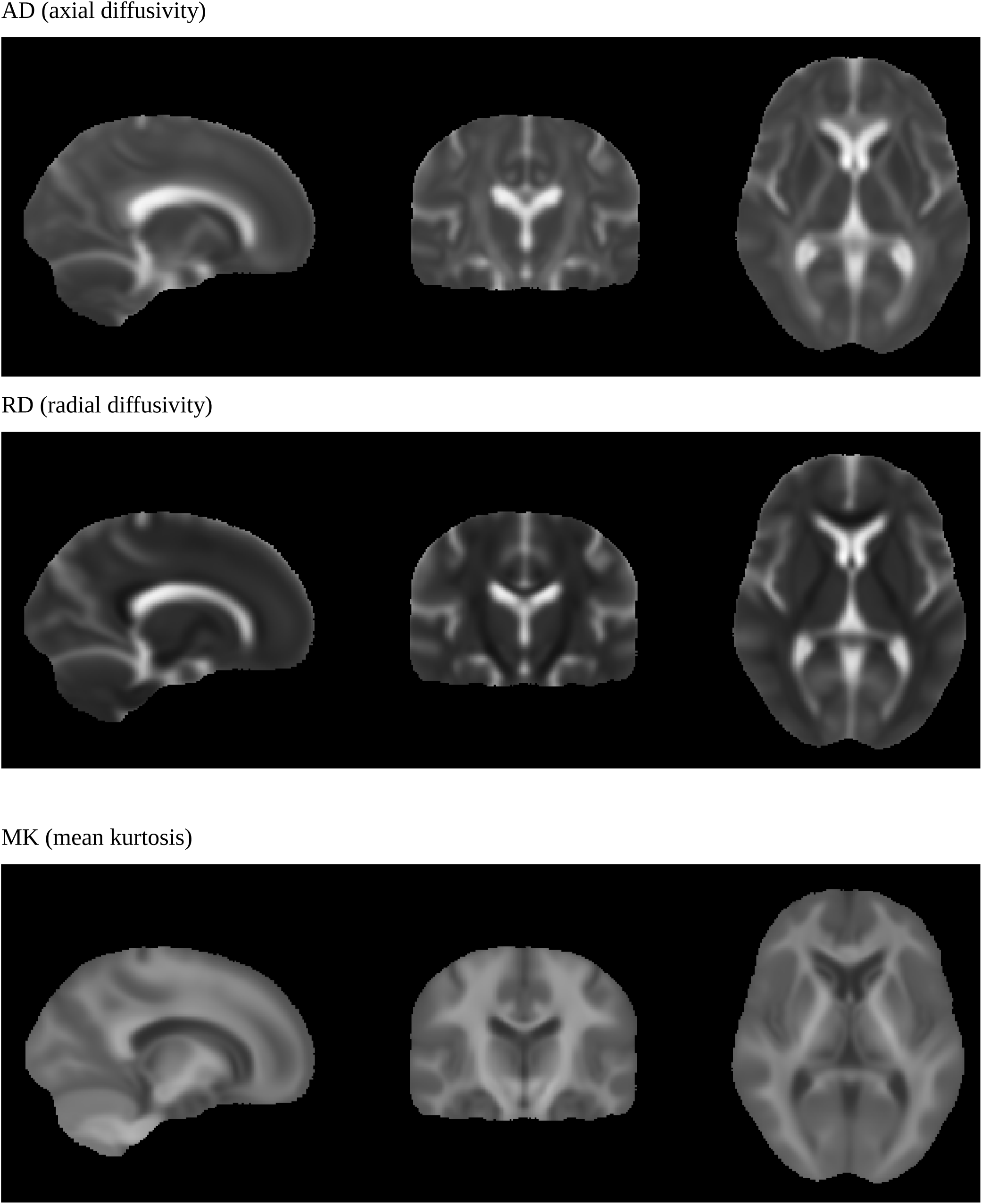

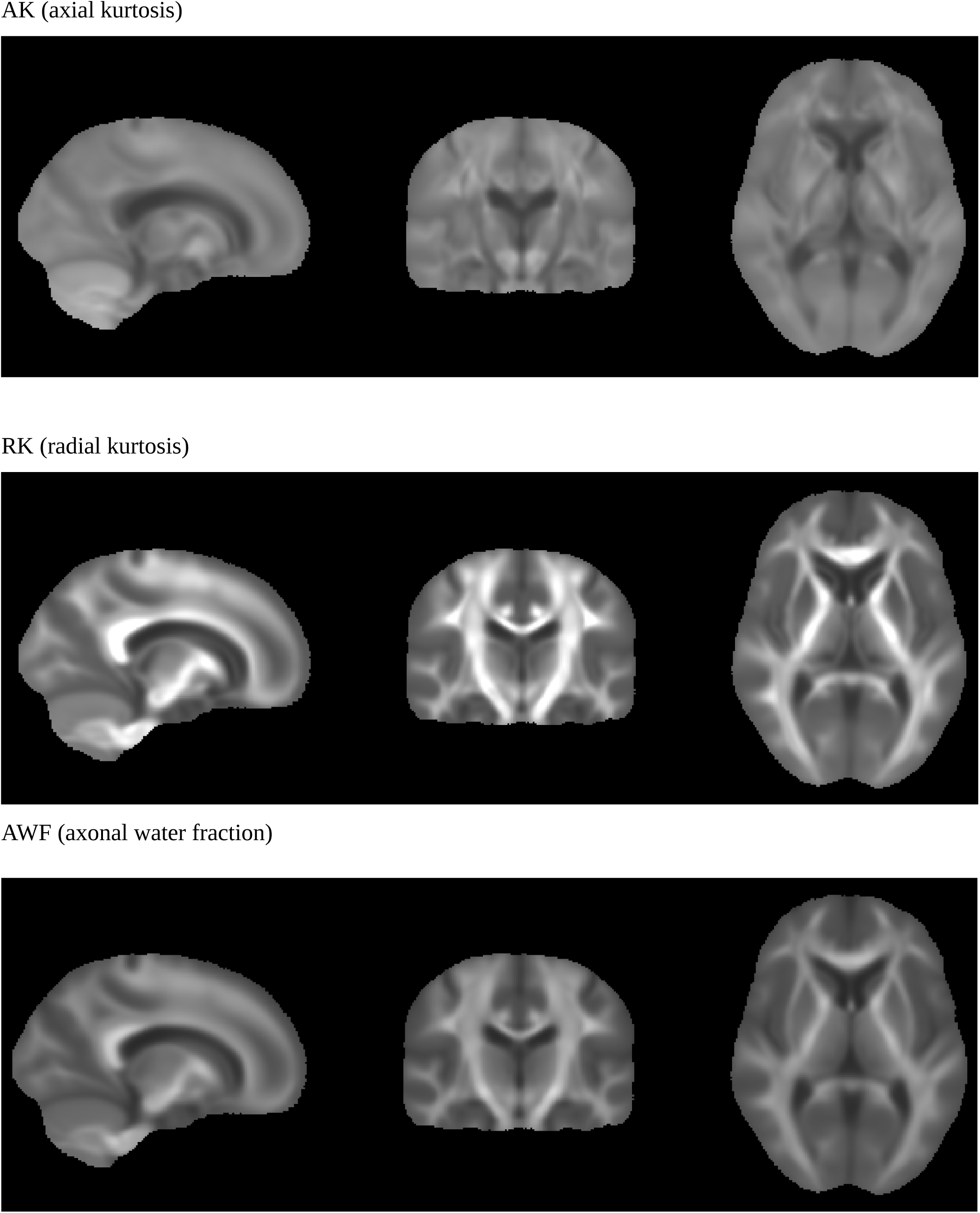

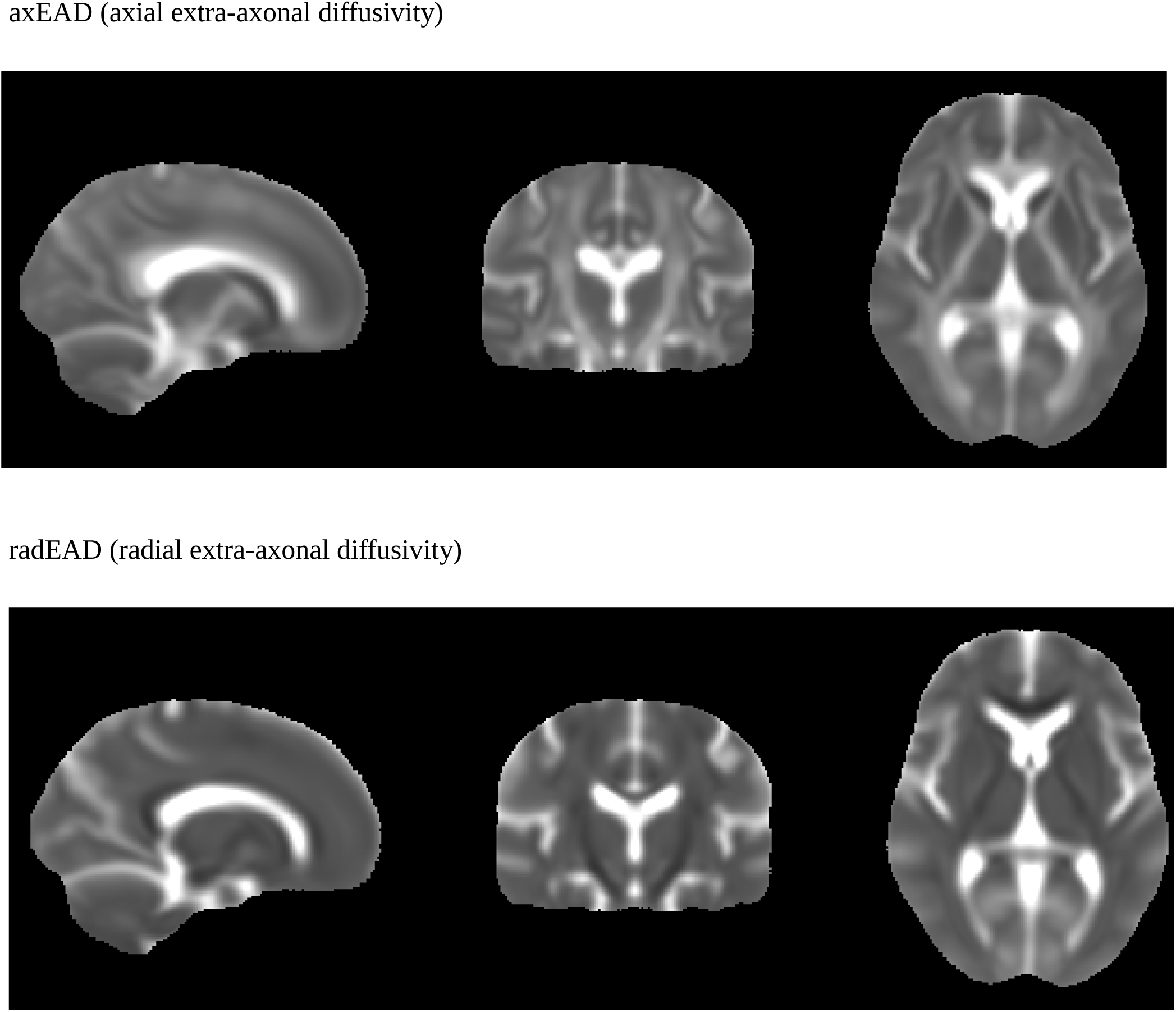

## 2. Age-diffusion correlations

The results of GLM models with linear and quadratic Age terms (black line) and with only linear Age term (red line). The diffusion data were QC filtered by the proposed method.

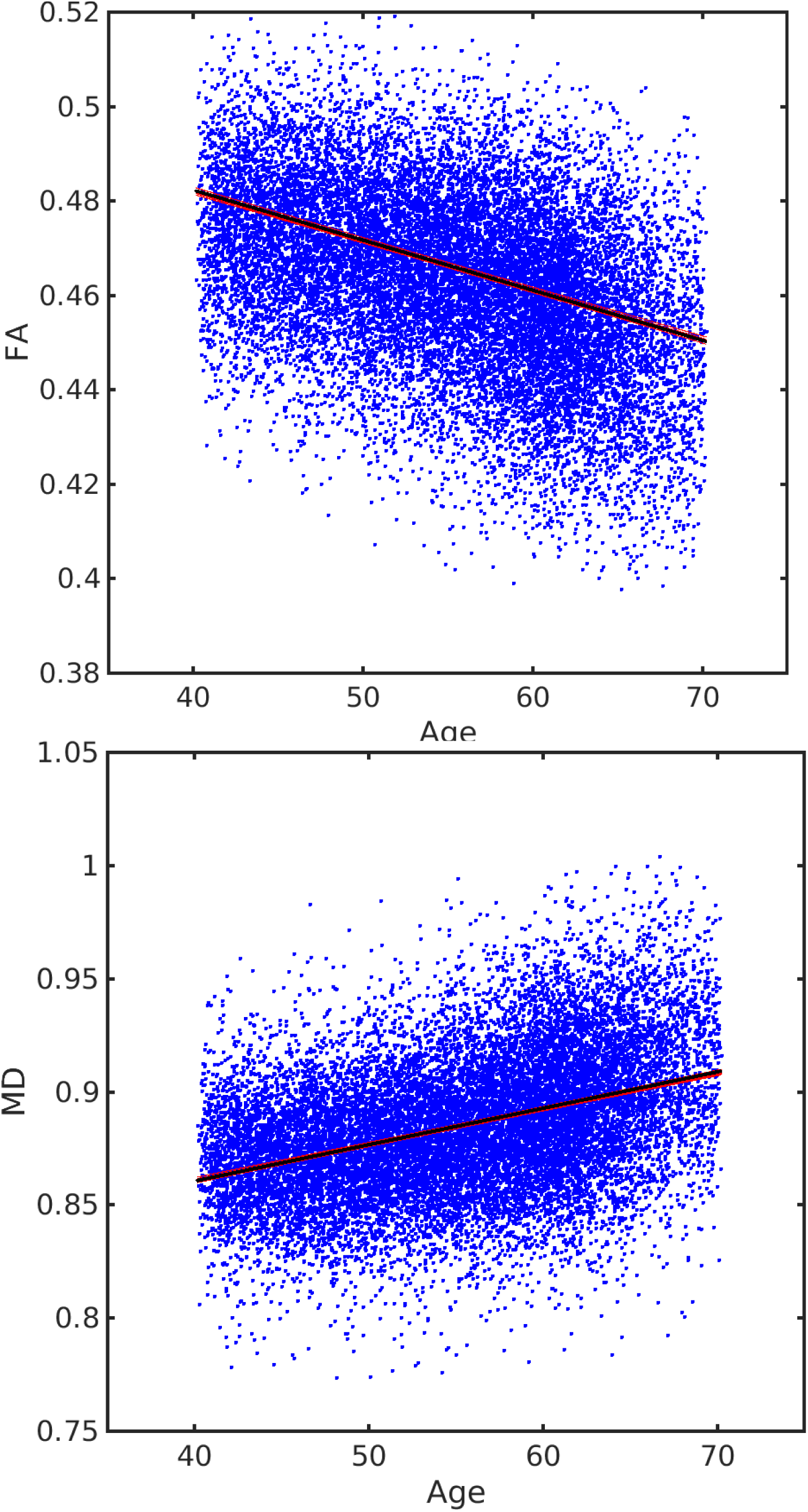

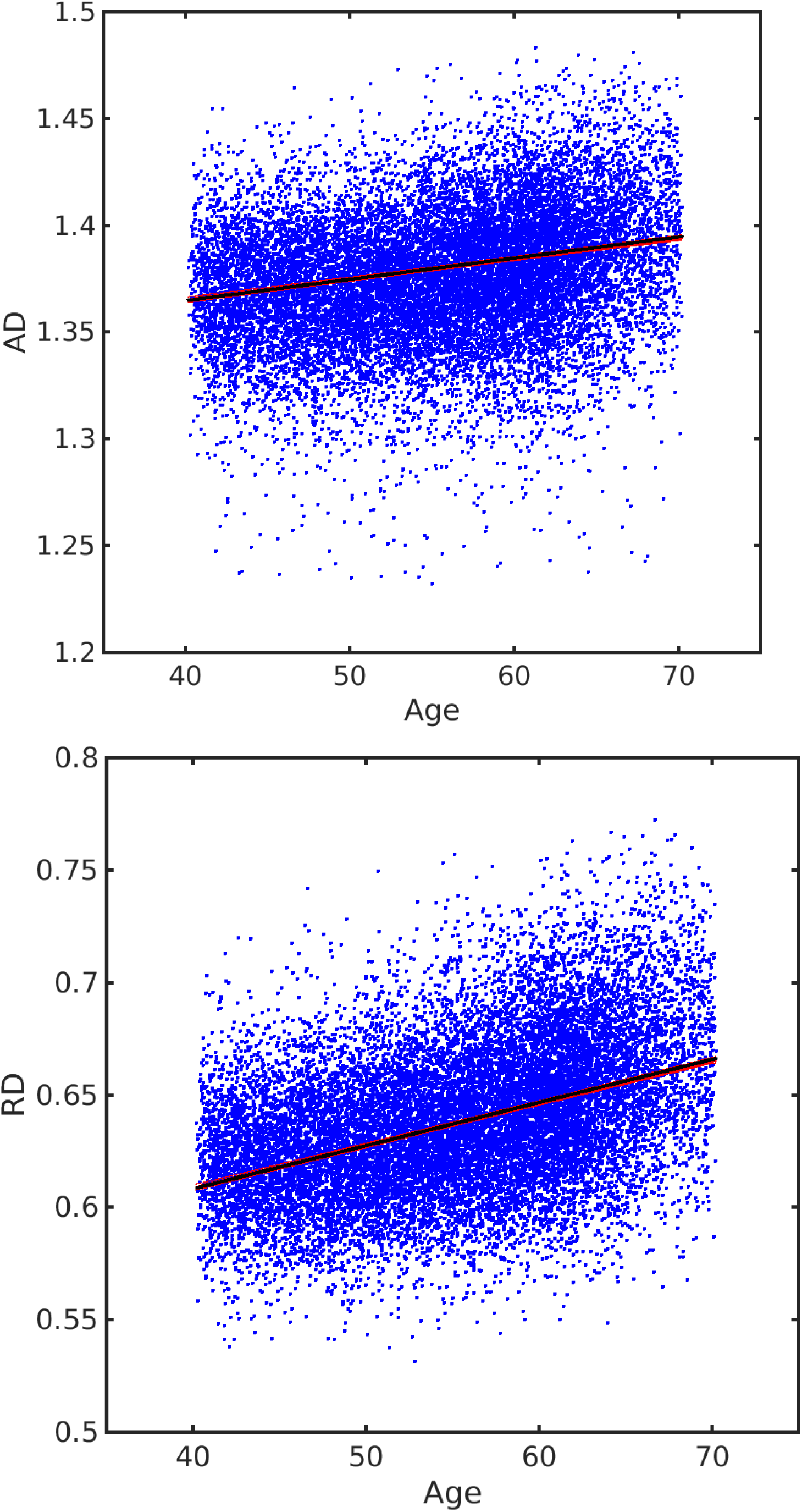

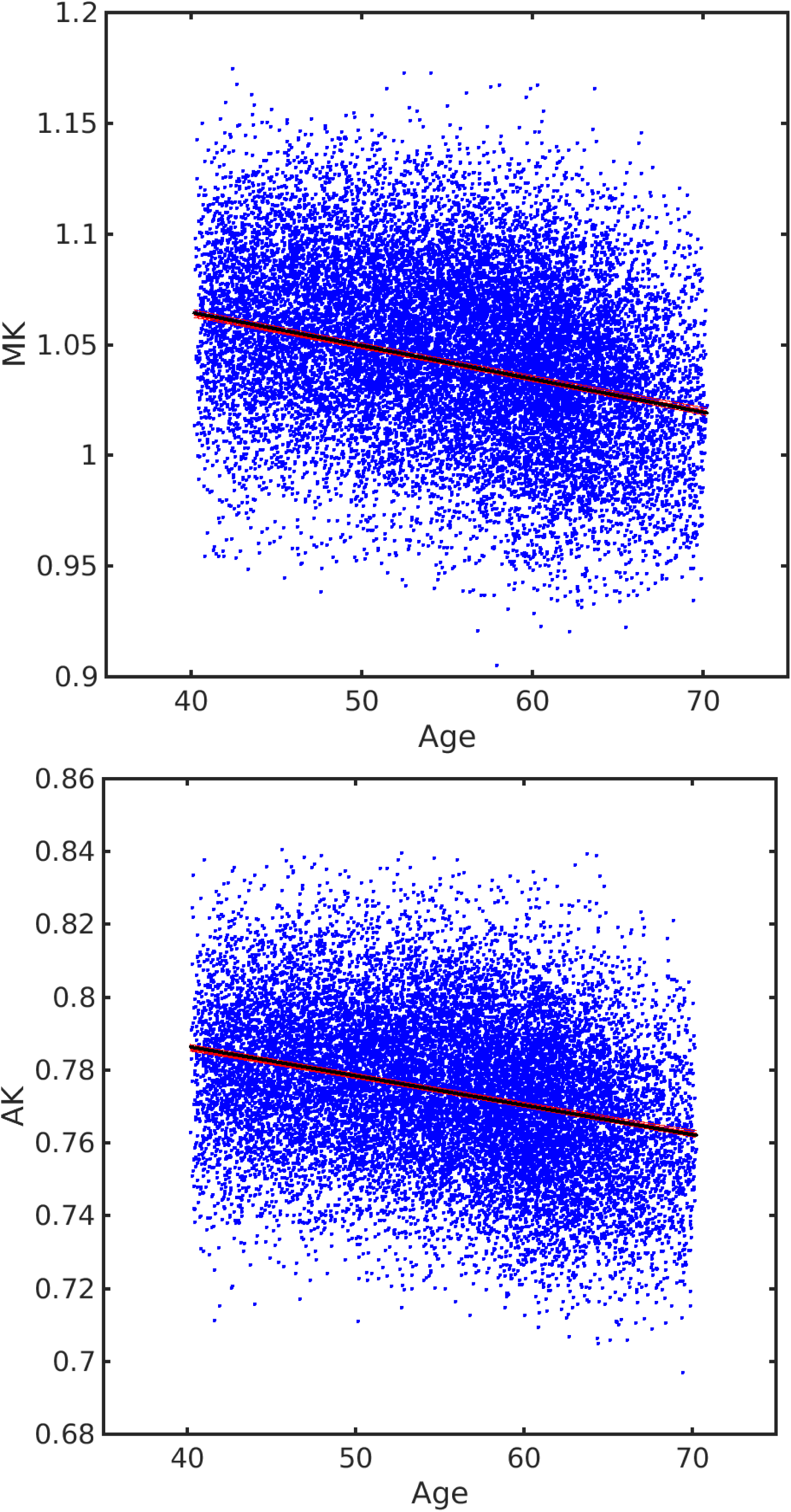

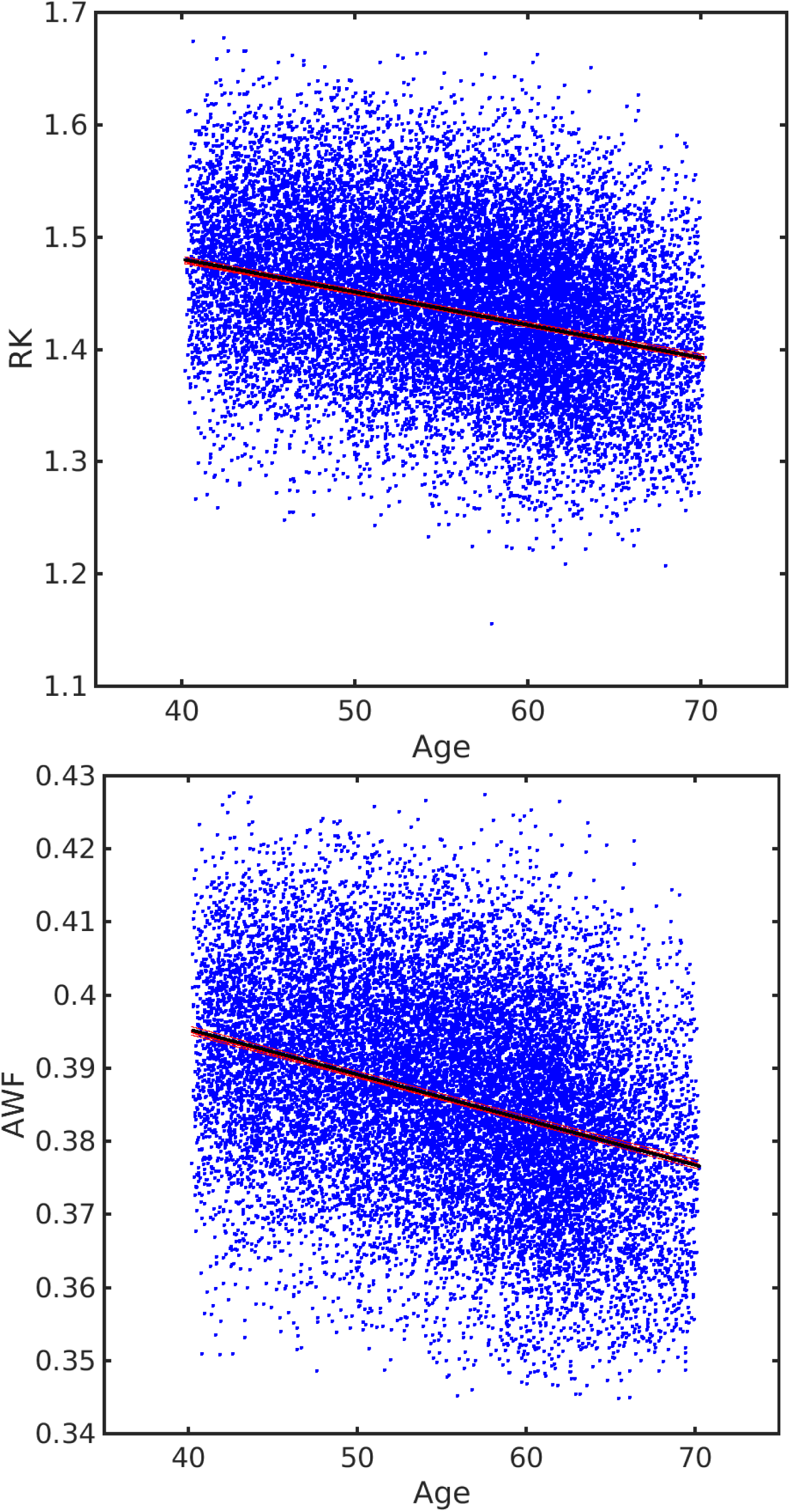

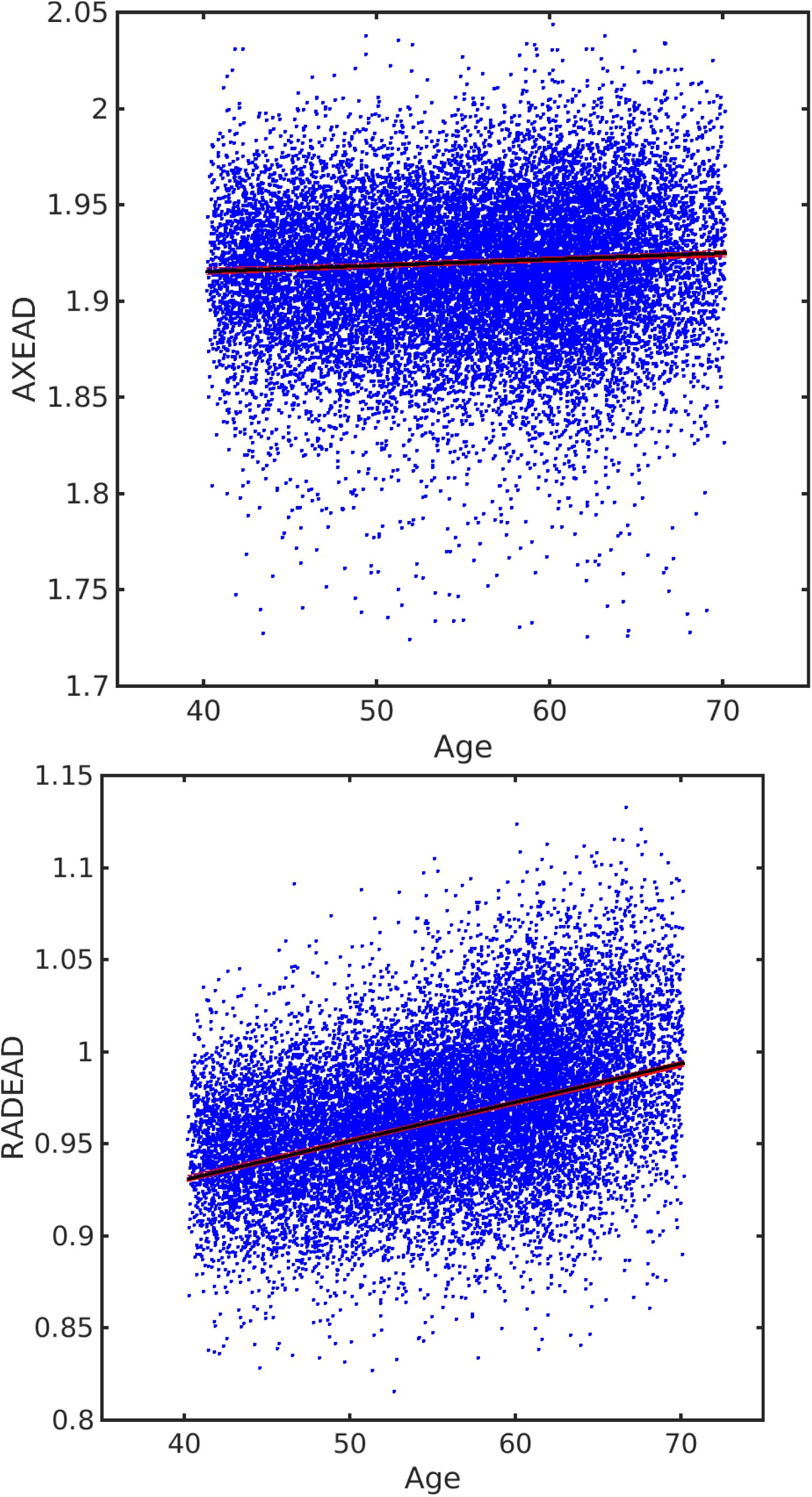

## 3. QQ plots of GLM model residuals with raw and QC filtered data

QQ plots are plotted for two datasets: original raw and QC filtered data. GLM models consists of two approaches with linear and quadratic Age terms. FA

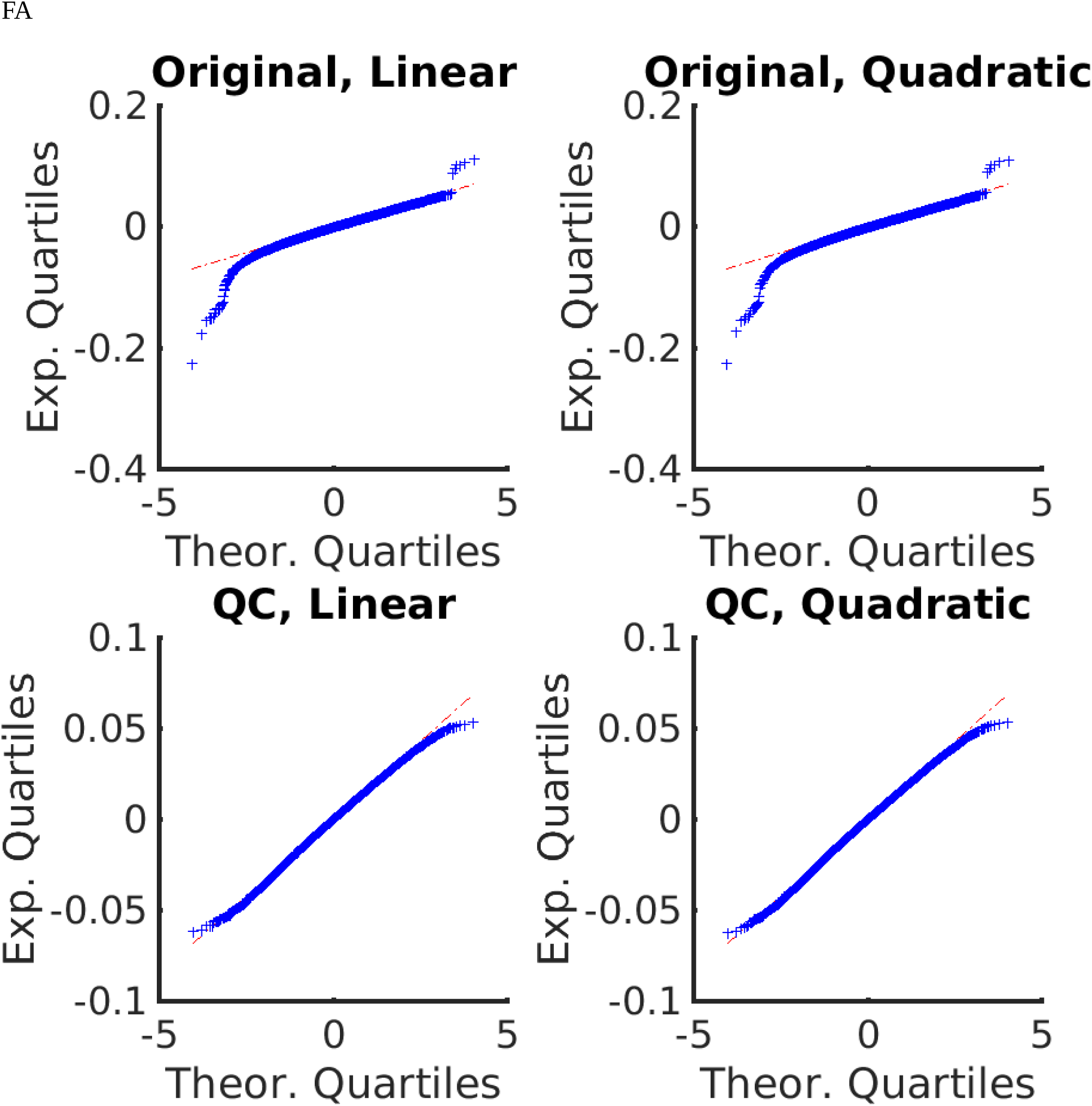

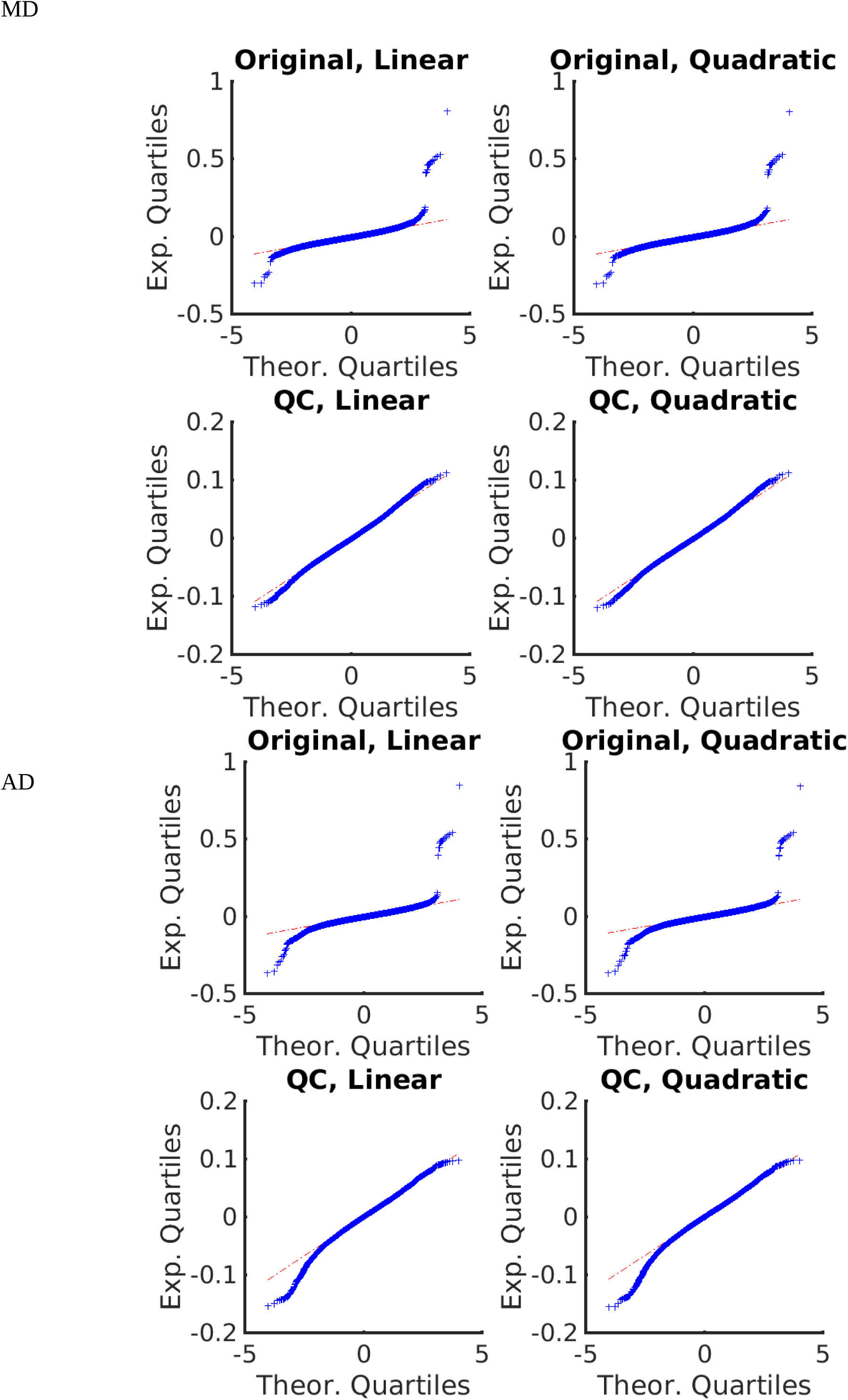

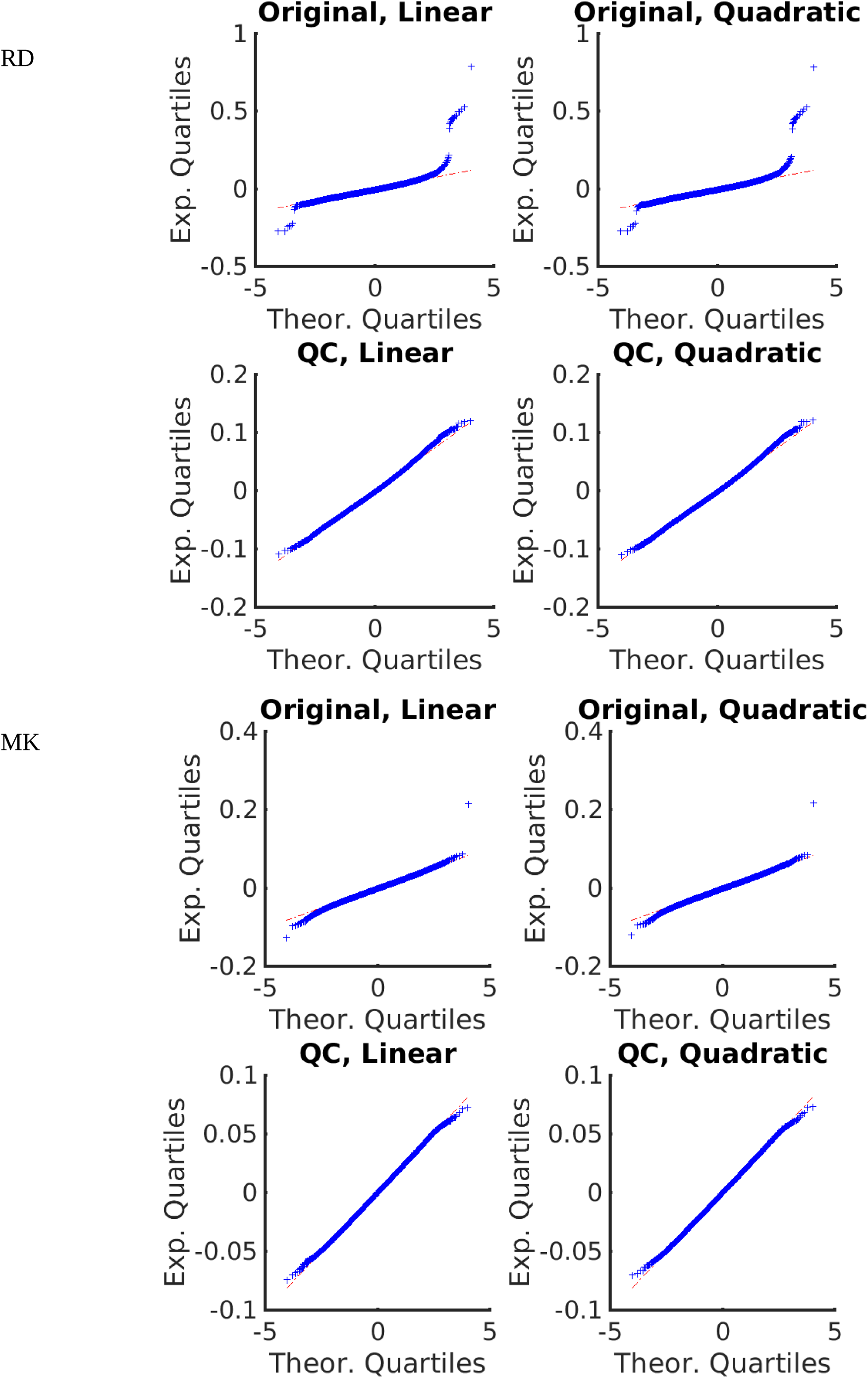

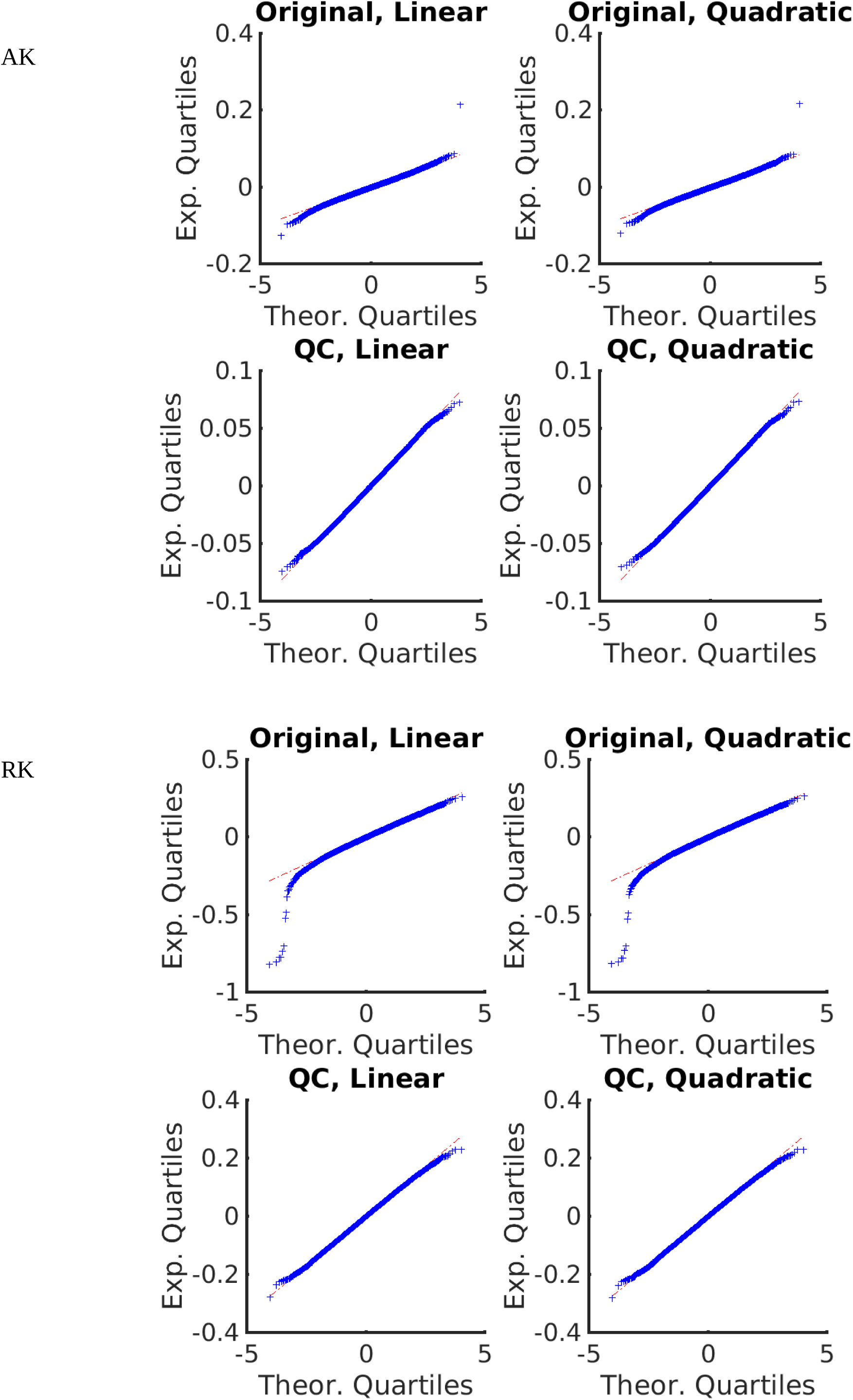

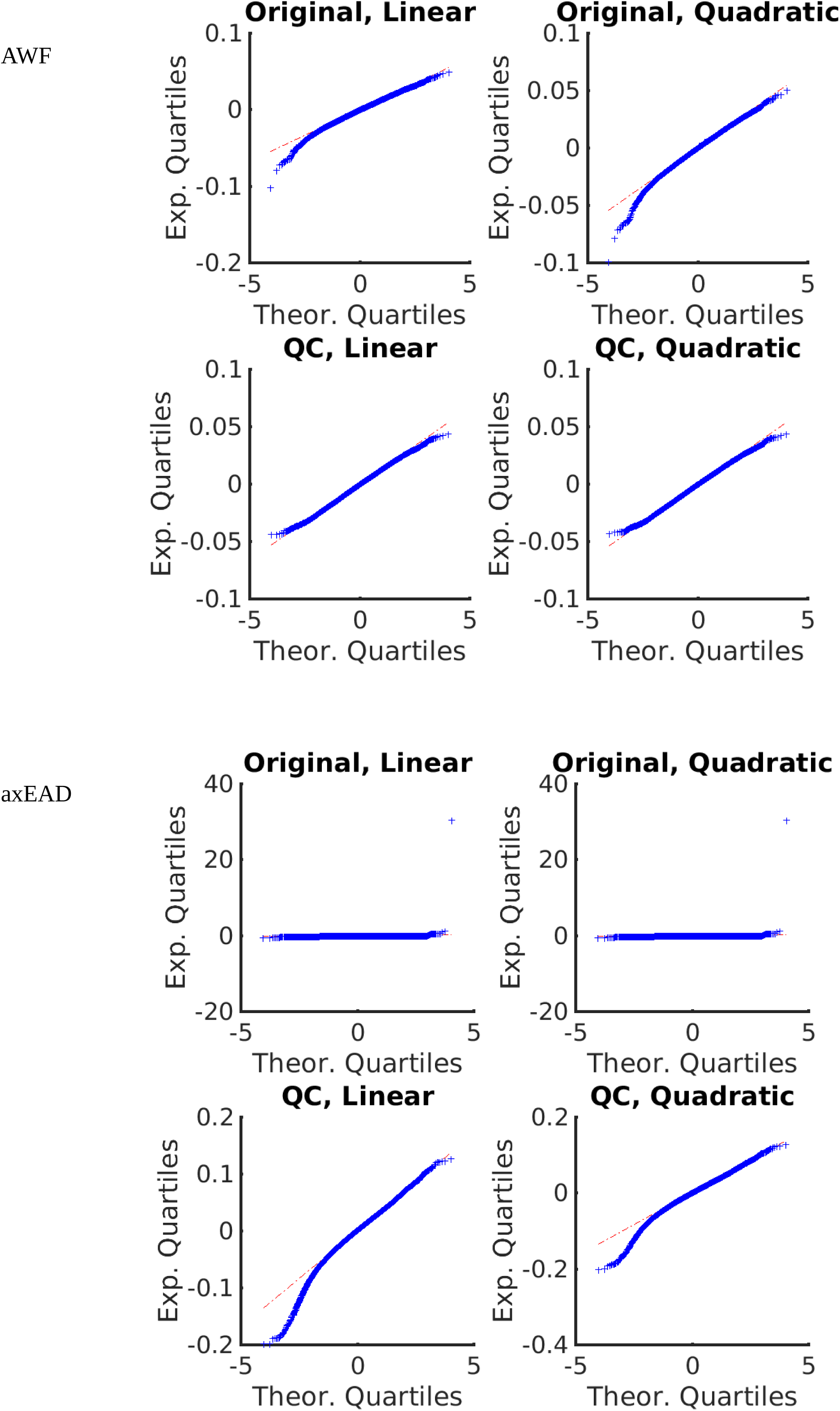

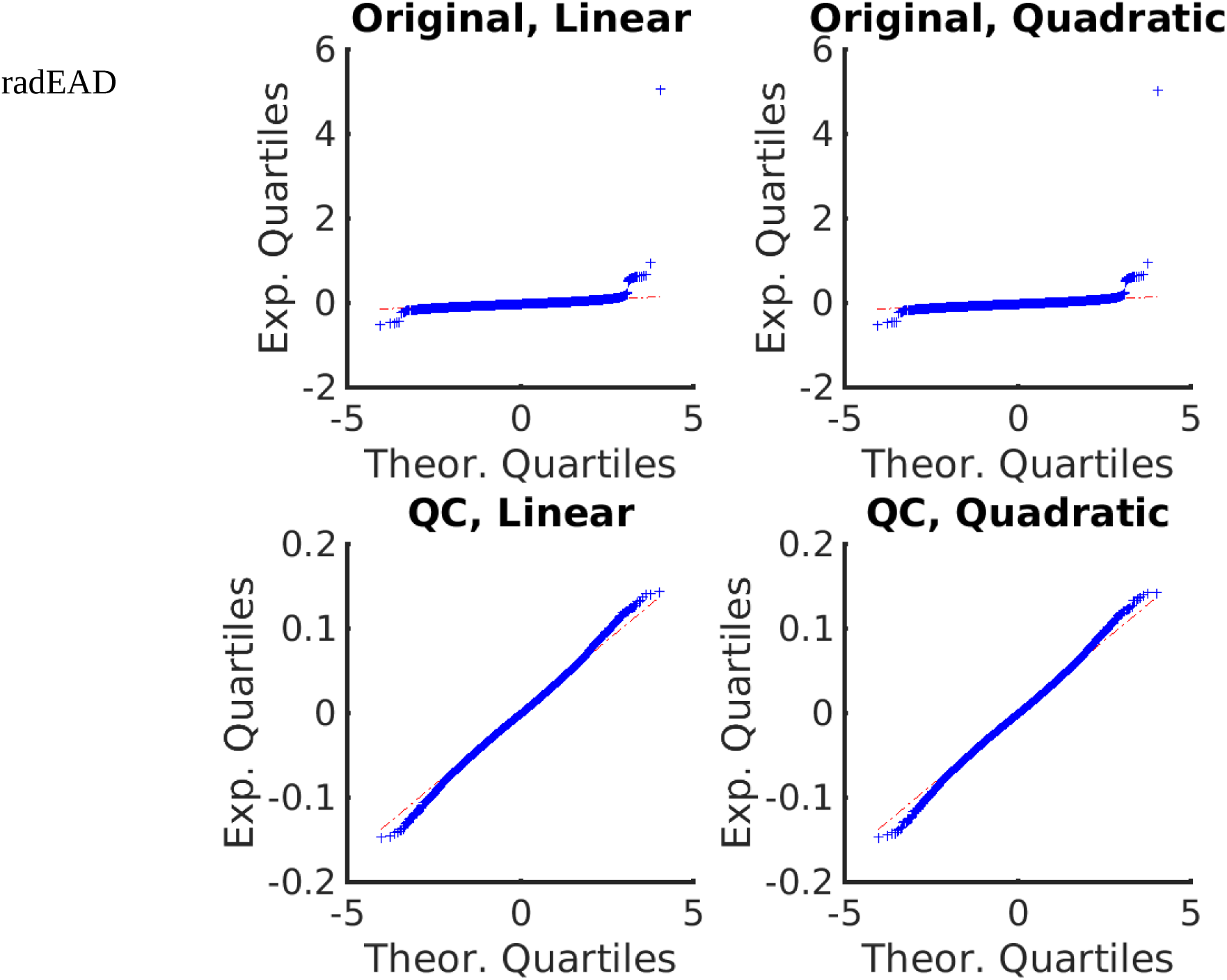

## 4. Demographic data of the detected outliers in UKB sample

**Figure.**
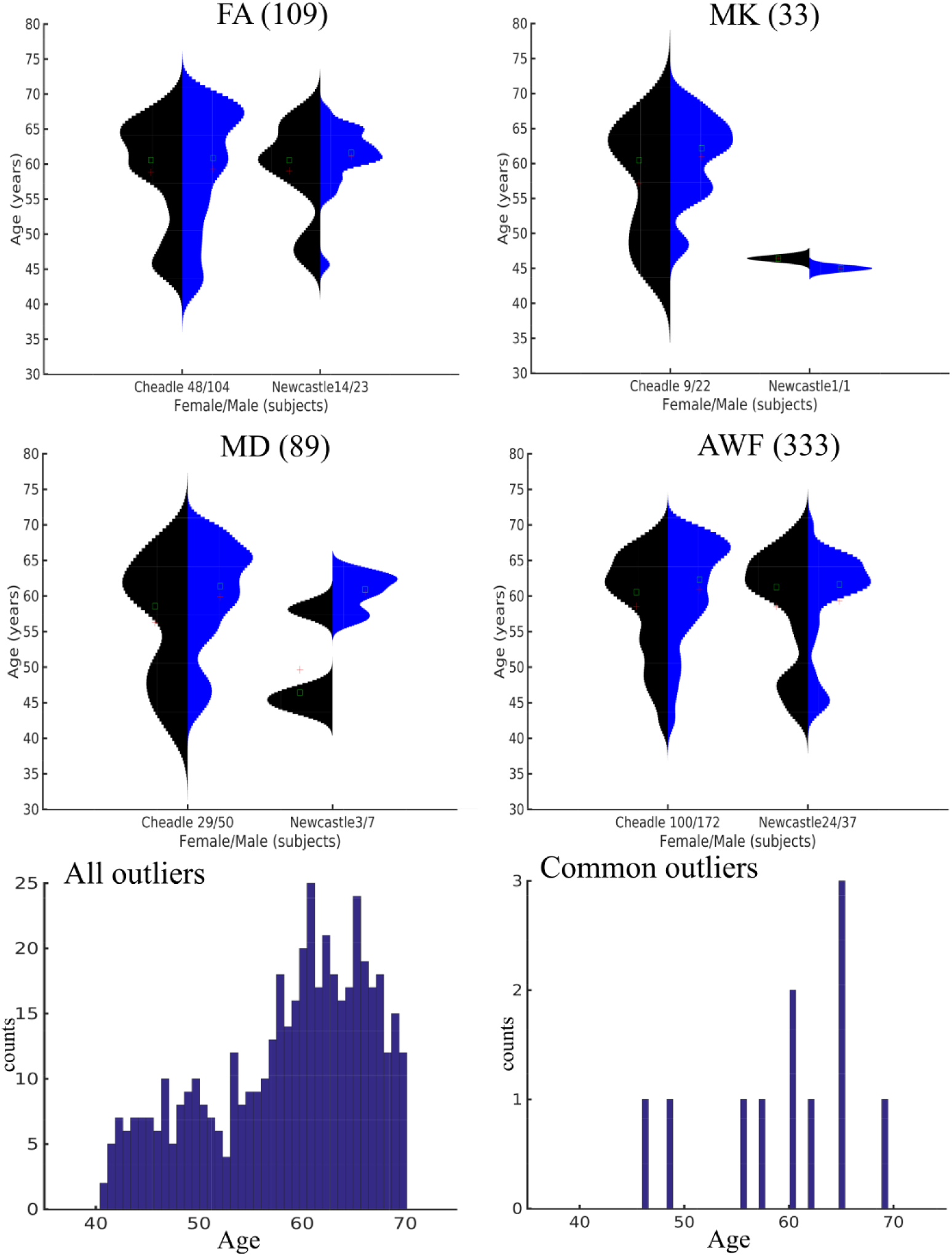
Demographic data of outlier-marked subjects detected over FA, MD, MK, and AWF metrics, respectively, separated into gender and scanner site groups. The bottom row plots present the age histograms for all outlier-marked subjects for DTI, DKI, and WMTI metrics together (left-side plot) and for the outlier-marked subjects appeared in all diffusion metrics.

